# APOBEC3A deaminase catalyzes site-specific editing of transfer RNAs

**DOI:** 10.1101/2025.06.10.658825

**Authors:** K. Hoffa-Sobiech, J. Sikora, M. Ruszkowski, M. Starczak, D. Gackowski, A. Błaszczak, A. Mierzwa, M. Rakoczy, M. Marcinkowska-Swojak, I. Stolarek, P. Jackowiak, M. Figlerowicz, L. Budzko

**Affiliations:** Institute of Bioorganic Chemistry Polish Academy of Sciences, Noskowskiego 12/14, 61-704, Poznan, Poland; Department of Clinical Biochemistry, Faculty of Pharmacy, Collegium Medicum in Bydgoszcz, Nicolaus Copernicus University in Torun, Karlowicza 24, 85-092 Bydgoszcz, Poland; Department of Gamete and Embryo Biology, Institute of Animal Reproduction and Food Research, Polish Academy of Sciences, Olsztyn, Poland

## Abstract

APOBEC3A is a cytidine deaminase that plays a crucial role in innate immunity; however, it can also drive oncogenesis when dysregulated. While its DNA editing activity is well-studied, the impact of APOBEC3A on RNA has only recently gained attention. Previous studies revealed that APOBEC3A deaminates mRNA stem-loop structures, however, its activity on other RNA classes remains unexplored. Given its likely evolutionary origin from tRNA adenosine deaminases and the prevalence of stem-loop structures in tRNA, we investigated APOBEC3A’s activity on tRNAs. We found that *in vitro* APOBEC3A efficiently deaminates a large spectrum of tRNA isoacceptors, primarily at anticodon positions. To assess whether the editing sites identified *in vitro* can be detected in tumor tissues, we analyzed data from The Cancer Genome Atlas. We identified six editing sites present in numerous patient samples. Our results point to a possible impact of APOBEC3A on tRNA decoding capacity, with potential relevance to mistranslation and cancer development.

## Introduction

APOBEC3A (Apolipoprotein B mRNA Editing Enzyme, Catalytic Polypeptide-Like 3A; A3A) is a member of the AID/APOBEC family [1]. The latter includes zinc-dependent deaminases that convert deoxycytidine (dC)/cytidine (rC) to deoxyuridine (dU)/uridine (rU) in DNA and/or RNA chains [2]. In the human genome, there are 11 genes encoding AID/APOBEC proteins: *APOBEC1, APOBEC2, APOBEC4, AID*, and seven *APOBEC3* genes: *APOBEC3A, APOBEC3B, APOBEC3C, APOBEC3D, APOBEC3F, APOBEC3G,* and *APOBEC3H* [3]. The C>U conversion catalyzed by these enzymes alters nucleotide sequence, and thus, affects genetic information or DNA/RNA functions. Similarly, to adenosine deaminases acting on RNA (ADARs), AID/APOBECs originated from an ancestor of tRNA adenosine deaminases (Tad/ADATs), which edit adenine (A) to inosine (I) in tRNAs of eukaryotes and prokaryotes. All these enzymes share common core structural features such as a canonical deaminase structural motif HxEx_25-30_PCx_2-4_C, in which conserved Cys and His residues together with a water molecule coordinate a zinc ion [1, 4]. The conserved catalytic center is surrounded by four amino acid loops L1, L3, L5, and L7, which are believed to be critical for substrate specificity [5, 6].

A3A is well known for its role in innate immunity by restricting viruses and mobile genetic elements like retrotransposons [7–10]. A3A targets single-stranded DNA (ssDNA) preferentially within the 5’TC motif, being the most active enzyme within the human AID/APOBEC family [11]. Recent reports suggest that DNA secondary structure can significantly influence A3A activity [12, 13]. For example, A3A displays higher activity on stem-loop than on linear ssDNA [14].

As a very potent DNA mutator, A3A can also be implicated in cancer development and may drive oncogenesis [15–17]. It has been shown that *A3A* is upregulated in a variety of cancer types, including head and neck, lung, and breast cancers, and its expression is associated with the accumulation of somatic mutations and epigenetic changes [18–20]. A3A is responsible for most mutations in human cancer attributed to single base substitution (SBS) signatures 2 and 13, although other family members (especially APOBEC3B) may also contribute [16].

So far, little is known about A3A involvement in RNA editing. It was reported for the first time by Sharma and coworkers, who demonstrated that A3A catalyzes site-specific C>U conversions in hundreds of mRNA transcripts in macrophages and monocytes [21]. Later studies confirmed this observation and showed that A3A targets regions capable of adopting stem-loop structures and preferentially deaminates C located in loops within the 5’UC sequence motif. The size of the loops, nucleotides flanking deaminated C, and stability of the stems have a major impact on the level of RNA editing [22]. Recently, Jalili and coworkers analyzed samples deposited in The Cancer Genome Atlas (TCGA) and confirmed the mRNA-editing in tumor cells in which the A3A was upregulated [23]. Although the mechanisms by which A3A recognizes and selectively edits RNAs remain unknown, one can speculate that this process may be controlled by a variety of factors, including the RNA structure, transcript accumulation level, post-translational modifications, and interactions with other proteins [21, 22, 24, 25].

RNA editing occurs not only in mRNAs but also in non-coding RNAs (ncRNAs), such as transfer RNAs (tRNAs) [26]. The most studied example of eukaryotic tRNA editing is ADAT-catalyzed A>I transformation that occurs at the wobble position and expands the codon recognition capability [27]. C>U-edited tRNAs have been identified in various organisms (Figure 1) [28–31]. Products of C deamination have been observed in three positions of the anticodon loop: 32, 34 (the wobble position), and 35 [29–34]. Moreover, C>U editing outside the anticodon loop (positions: 4, 6, 8, 12, 28, and 41) has been postulated as a mechanism increasing the stability of tRNA molecules [28, 35–37]. The enzymes responsible for the tRNA editing remain largely unknown. The exceptions are CDAT8 and ADAT2/ADAT3, most likely involved in the edition of positions 8 [28] and 32, respectively [33, 38].

**Figure 1.**
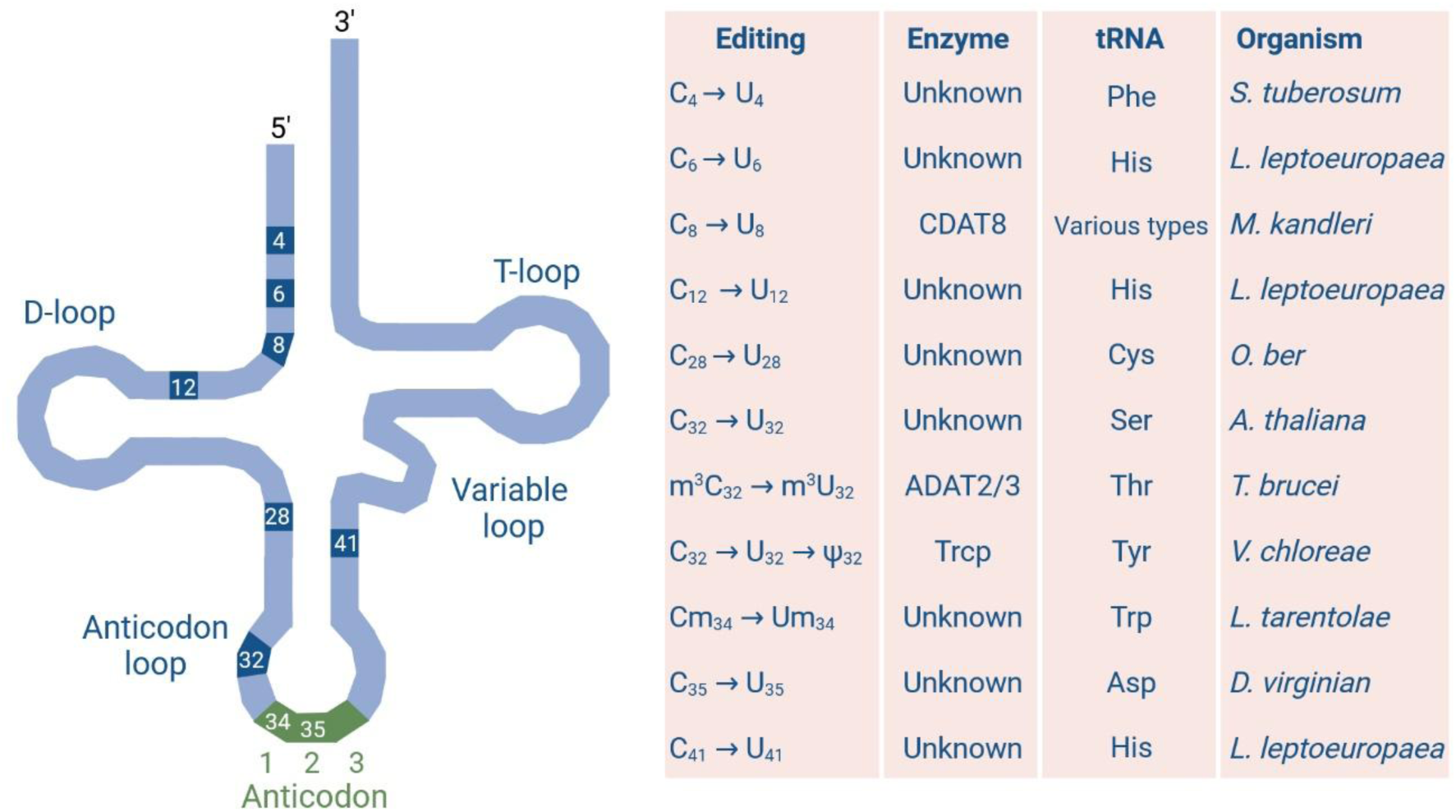
Reported examples of C>U editing in tRNAs. Positions 34, 35, and 36 usually (depending on the length of tRNA) correspond to the first (the wobble), second, and third anticodon positions, respectively. Created in BioRender. Hoffa-Sobiech, K. (2025) https://BioRender.com/lh61li4.

There are many reports demonstrating that mature tRNAs may undergo enzymatic cleavage (mostly within loops) into shorter derivatives. Some of them were shown to accumulate in cells and function as a novel class of small regulatory RNAs. Because the classification and nomenclature of these molecules are complicated [39], we will collectively call all of them tRNA-derived fragments (tRFs). There is increasing evidence that tRFs are involved in various molecular processes, including but not limited to gene silencing, transcription regulation, and translation control [39]. tRFs are also involved in many pathophysiological processes, including cancer, where a correlation between tumor progression and tRFs accumulation has been demonstrated [40]. Despite C>U deamination has been observed in tRNAs, the field of tRF editing remains practically unexplored.

Considering that: (i) AID/APOBEC family most likely evolved from tRNA adenosine deaminases, (ii) A3A edits RNA capable of adopting stem-loop structures and (iii) within a stem-loop structure A3A preferentially deaminates C located in loop within 5’UC motifs, we hypothesized that A3A might be the key enzyme participating in tRNA editing in humans. To verify this assumption, we tested A3A activity on tRNA in vitro and then searched for A3A-specific tRNA editing in cancer samples. We showed that recombinant A3A edits *in vitro* both cell-derived and synthetic tRNAs. We identified efficient editing of 17 mature tRNA isoacceptors. We found that the second and third positions of the anticodon were edited the most frequently and the most efficiently. However, we also observed editing at the first anticodon position, at position 38, as well as within variable and D loops. In synthetic tRNAs, we observed editing almost exclusively at the third anticodon position. By applying *in silico* modeling and molecular dynamics simulations, we proposed a mechanism of anticodon nucleobases recognition by A3A. Next, to assess whether the editing phenomenon can be observed in cancer, we compared miRNA-Seq, RNA-Seq, and WGS data obtained for individual samples. We found six putative A3A editing sites in tRFs. They were most frequently observed in thyroid cancer (THCA), lung adenocarcinoma (LUAD), head and neck squamous cell carcinoma (HNSC), BRCA-related cancer (BRCA), and ovarian cancer (OV). In general, our findings on A3A-mediated tRNA editing open up new perspectives in the studies of the mistranslation phenomenon, cancer development, and host defense against viral infections. Since A3A has been currently utilized in CRISPR-mediated RNA base editing tools [41], our discovery is also of significant practical importance. It may also have potential applications in developing new antiviral therapies and detecting new tumor markers.

## Results

### Recombinant A3A edits *in vitro* tRNAs isolated from HEK293T cells

To test A3A activity on tRNA, we produced recombinant wild-type enzyme (WT_A3A) and its catalytically inactive mutant E72A (E72A_A3A) in a bacterial expression system, as earlier described [42, 43]. Both variants were purified (Supplementary Figure S1), and their activities on canonical substrates were confirmed by *in vitro* ssDNA and RNA deamination assays. The reaction products were analyzed by Ultra Performance Liquid Chromatography (UPLC). As the canonical substrates, we used synthetic 80-nucleotide ssDNA (containing a single C residue within a 5’TC motif; ssDNA_80nt) and a 14-nucleotide RNA, being the fragment of the succinate dehydrogenase B gene transcript (SDHB_14nt). The latter has been shown to form a stem-loop structure and is the most studied substrate for A3A activity on RNA [21, 22, 44]. Both ssDNA_80nt and SDHB_14nt were incubated with WT_A3A or E72A_A3A, and reaction products were digested to single nucleosides, followed by UPLC analysis. As shown in Figure 2 and Supplementary Figure S2, we observed high activity of WT_A3A on both canonical substrates. Deamination of ssDNA_80nt resulted in a decrease of the dC content in the total pool of nucleosides by almost 100% (Supplementary Figure S2, panel B). In the case of the SDHB_14nt substrate, only one of the four Cs was available for deamination. Therefore, we observed a subtler decrease in the total rC content (from 30.2% to 22.6% in the E72A_A3A and WT A3A-treated samples, respectively). Such a result corresponds to a 100% decrease of rC content if only one C position available for deamination is considered (Supplementary Figure S2, panel C). The observed changes were accompanied by a corresponding increase in the content of dC/rC deamination product, i.e. dU/rU (Figure 2A and 2B).

**Figure 2.**
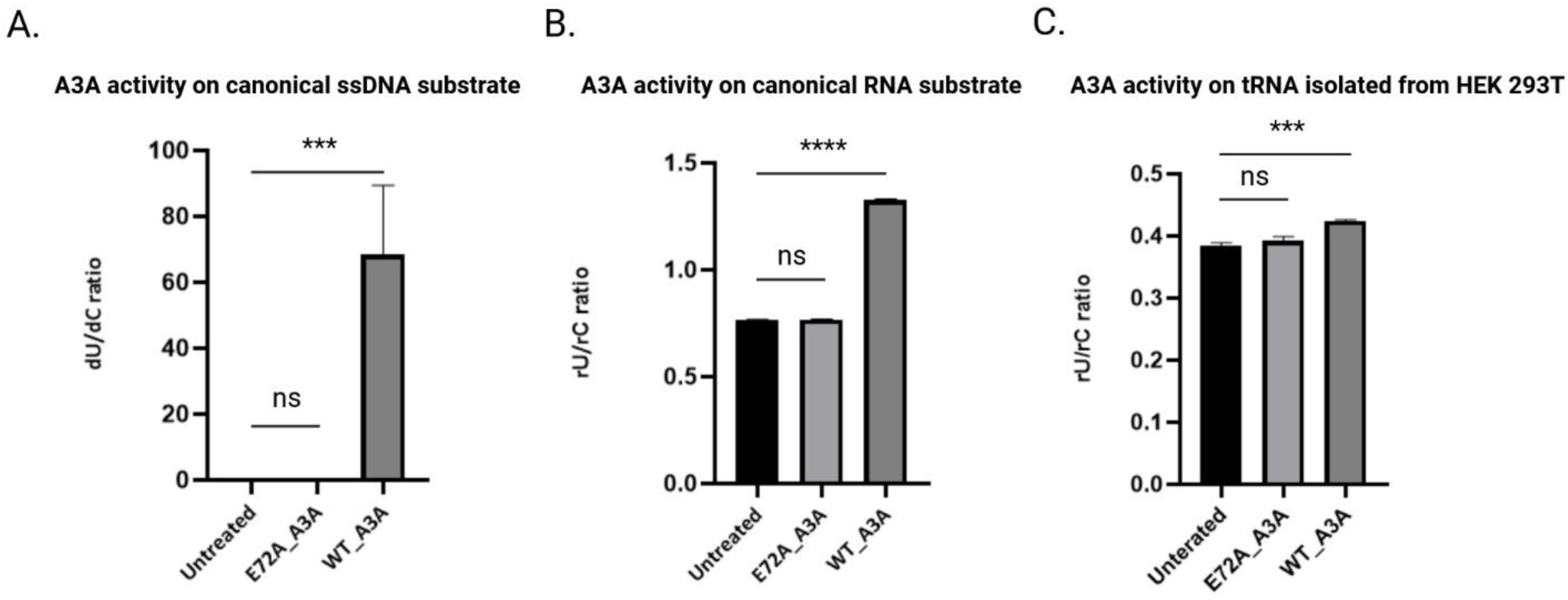
Quantitative analysis of A3A activity on canonical DNA and RNA substrates, and cell-derived tRNA. A3A activity on ssDNA_80nt (A), SDHB_14nt (B), and tRNA isolated from HEK293T cells (C). WT_A3A- or E72A_A3A-treated and untreated substrates were digested to nucleosides and analyzed by UPLC. The deaminase activity was expressed as a product-to-substrate (rU/rC) ratio (average of three replicates). P-values were calculated with ordinary one-way ANOVA and Dunnett’s multiple comparisons test. ***p <0.001, ∗∗∗∗p <0.0001. Created in BioRender. Hoffa-Sobiech, K. (2025) https://BioRender.com/hosin3z.

To determine whether WT_A3A catalyzes the conversion of C>U in native tRNAs, the fraction containing these molecules was isolated from HEK293T cells. Then, the assay described above involving recombinant WT_A3A or E72A_A3A was performed. As earlier, deamination products were detected by UPLC. The high quality of the isolated tRNA fraction was confirmed using the Agilent TapeStation System (Supplementary Figure S3). As shown in Supplementary Figure S2 (panel D), in the WT_A3A-treated sample, we observed a 1% decrease of the rC content in the total pool of nucleosides, as compared to the control samples treated with E72A_A3A or untreated. Such a decrease in rC content means that in one in four tRNA molecules, one C has been edited (assuming that the average length of a tRNA is 83 nucleotides and the average rC content is 32%). The above was confirmed by the increase of rU to rC concentration ratio in the WT_A3A-treated sample, compared to the E72A_A3A-treated or untreated control samples (Figure 2C). The statistical significance of our result was confirmed in three replicates.

### Structural requirements of A3A-mediated tRNA editing

In the next step, we tried to determine if there is some pattern of tRNA editing by A3A or if it is a random process. To this end, we performed high-throughput sequencing of tRNA isolated from HEK293T cells and treated with WT_A3A. As negative controls, we used tRNA isolated from HEK293T and treated with E72A_A3A or not treated with any enzyme. To prepare NGS libraries, we applied the YAMAT-seq protocol [45], designed specifically for tRNA sequencing. As a result, we identified 87 nucleus-encoded tRNA isodecoders for 32 isoacceptors (Supplementary Table S1) having the required coverage (250x). The positions edited in tRNA were identified using a rigorous protocol that led us to identify 80 edited positions within 55 tRNA isodecoders (Supplementary Table S2). The editing efficiency for detected positions ranged from 1.6% to 98.8%. Importantly, the estimated background level of editing in both controls was negligible (on average ∼0.4% for E72A_A3A-treated and ∼0.2% for untreated samples). To better define the sequence/structure preferences of the A3A activity, we further restricted our analysis to nucleotides edited with at least 10% efficiency. We found 45 such nucleotides located within 35 tRNA isodecoders, corresponding to 17 isoacceptors (Table 1, Figure 3, and Figure 4). The highest editing efficiencies were observed for tRNA-Arg-TCG (98.8%), tRNA-Glu-TTC (98.5%), tRNA-Asp-GTC (98.3%), and tRNA-Arg-CCG (98.1%). Most isoacceptors (13 out of 17) were edited in the anticodon loop, mostly within the anticodon. The third and second anticodon positions were modified most frequently (the third position in 6 isoacceptors and the second position in 4 isoacceptors) and with the highest efficiency. More than 98% of editing efficiency was observed in the third position of tRNA-Asp-GTC and tRNA-Glu-TTC. Above 97% of editing efficiency was found in the second position of tRNA-Arg-CCG and tRNA-Arg-TCG. Notably, the detected A3A-mediated editing at the second and third anticodon positions may expand the decoding capacity (see Discussion). The first anticodon position was edited in 4 isoacceptors but with lower editing efficiency (e.g., ∼73% in tRNA-Lys-CTT and ∼51% in tRNA-Met-CAT). In three isoacceptors, the anticodon loop was edited out of the anticodon at the loop’s 3’ end (position 38), however, with relatively low efficiency. Maximal editing efficiency at this position was detected for tRNA-His-GTC and reached 19.5%. For several tRNA isoacceptors, we observed A3A-mediated editing outside the anticodon loop. tRNA-Asn-GTT and tRNA-Lys-CTT were deaminated within the D-loop (position 17) with ∼10% efficiency. tRNA-Asn-GTT was edited solely within the D-loop. Four tRNA isoacceptors (tRNA-Leu-TAA, tRNA-Leu-TAG, tRNA-Ser-AGA, tRNA-Sec-TCA) were edited within the variable loop, and three of them (tRNA-Leu-TAA, tRNA-Leu-TAG, tRNA-Ser-AGA) were solely edited within this loop and were not edited in the anticodon loop. For tRNA-Sec-TCA and tRNA-Leu-TAG, the variable loop editing was very effective, reaching ∼85% and ∼73%, respectively. Notably, editing within the variable loop was observed only in tRNAs that contain an extended variable region. Similarly, to the anticodon loop, editing occurred in the middle of the variable loop (tRNA-Leu-TAA, tRNA-Leu-TAG, tRNA-Ser-AGA) or at its 3΄ end (tRNA-Sec-TCA). Interestingly, all isoacceptors edited exclusively outside the anticodon loop lacked C in the anticodon sequence. Six isoacceptors were edited in two positions. As shown by the WebLogo representation in Figure 4B, the edited Cs predominantly occurred in the 5’YU<u>C</u>R (Y=U/C, R=A/G) sequence motif, where deaminated C is underlined. An interesting example was tRNA-Sec-TCA, in which position 56 within the variable loop, within the unfavorable 5’GC context, was more effectively edited than the second anticodon position within the favorable 5’UC context. This suggests that other structural factors than sequence context may also significantly influence editing efficiency.

**Figure 3.**
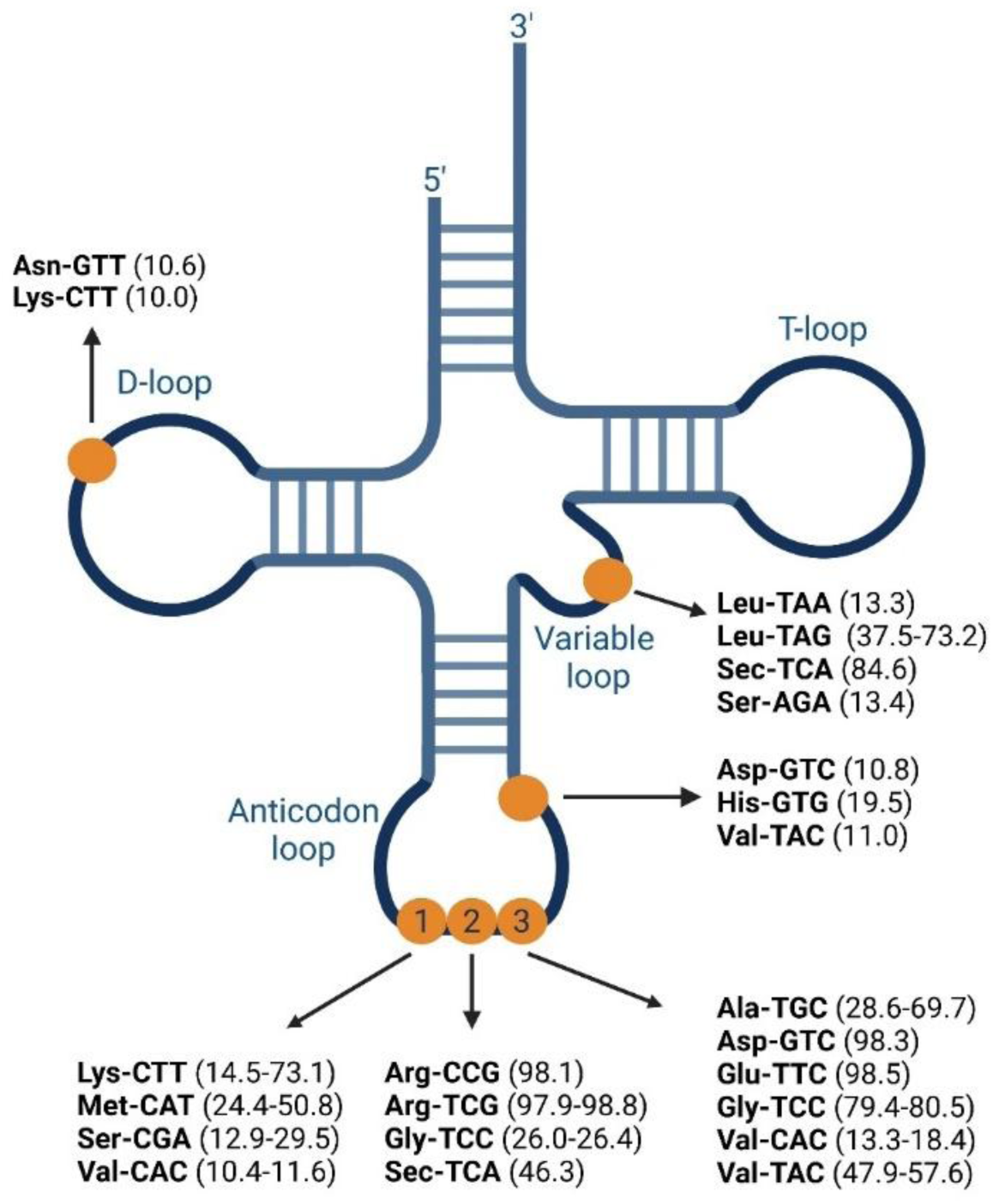
Schematic representation of C>U edited positions identified in tRNA isoacceptors. All positions edited with efficiency above 10% are shown. The orange dots represent deaminated Cs. The editing efficiency (in %) is shown in parentheses (a range is shown if edited positions were mapped to more than one isodecoder). Created in BioRender. Hoffa-Sobiech, K. (2025) https://BioRender.com/qd9gxk0.

**Figure 4.**
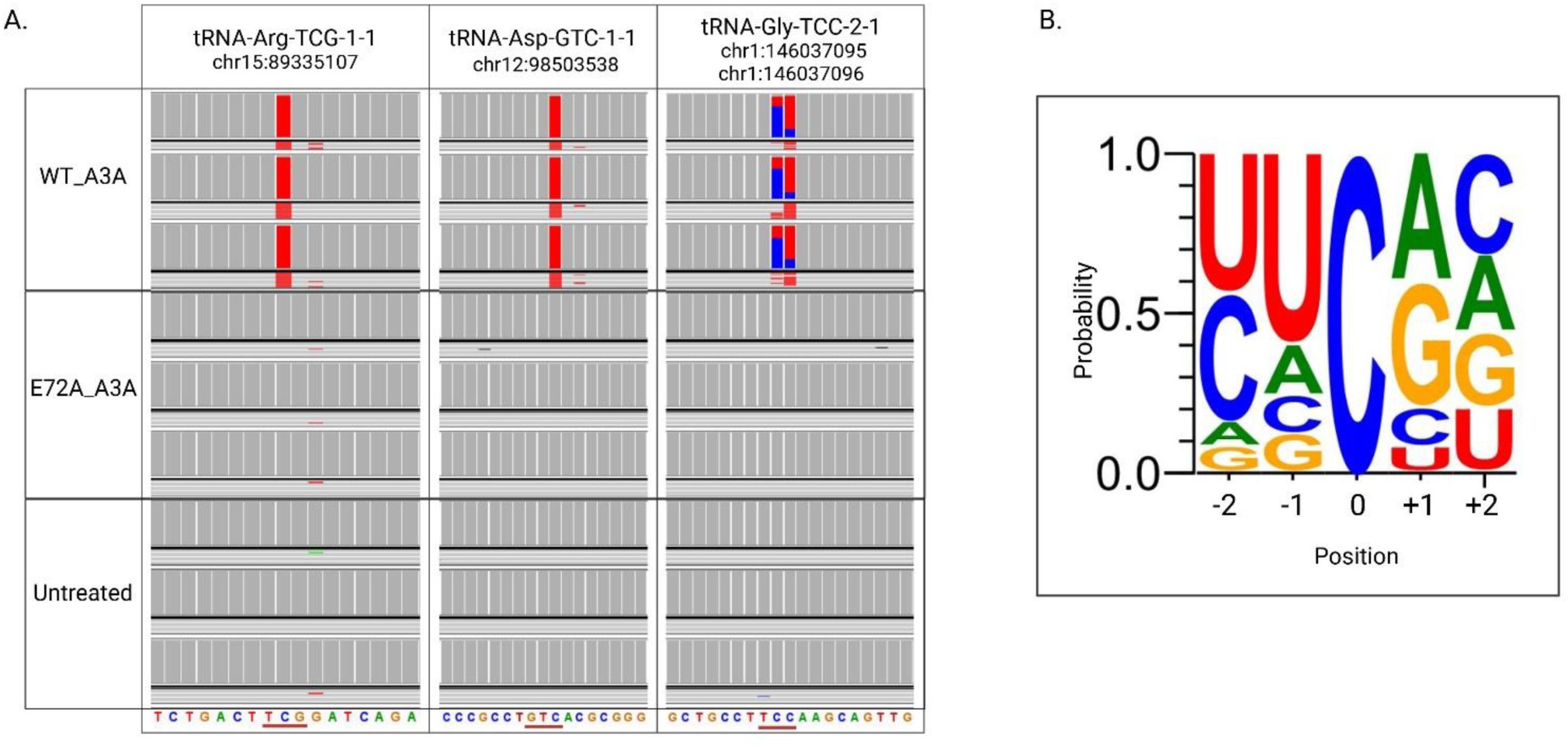
C>U editing within anticodon loops of tRNA isodecoders. (A) tRNA sequencing reads mapped to their genes (visualized with Integrative Genomics Viewer (IGV)). Alignments were generated for tRNA treated with WT_A3A and corresponding controls (E72A_A3A-treated and untreated samples). The reference sequence is shown at the bottom of each alignment, and the anticodon is underlined. Gray indicates no change compared to the reference sequence. C>T mismatches are presented as a proportion of red (for T) and blue (for C) colors corresponding to the percentage of editing. (B) WebLogo representation of the local sequences around the C (±2 nucleotides from C) efficiently edited by WT_A3A. Created in BioRender. Hoffa-sobiech, K. (2025) https://BioRender.com/yhjez2u.

**Table 1.** tRNA isoacceptors edited by WT_A3A in at least one position with efficiency higher than 10%. Mean editing efficiency, sequence motif within which deaminated C was located, and the tRNA regions where deamination occurred are indicated. If deamination occurs in the anticodon, the exact anticodon position is specified.

| Lp. | tRNA isoacceptor | Mean editing [%]* | Sequence motif** | tRNA region | Position in the anticodon |
| --- | --- | --- | --- | --- | --- |
| 1 | Ala-TGC | 28.6-69.7 | TG**C**AT | Anticodon loop | 3 |
| 2 | Arg-CCG | **98.1** | TCCGG | Anticodon loop | 2 |
| 3 | Arg-TCG | **97.9-98.8** | TTCGG | Anticodon loop | 2 |
| 4 | Asn-GTT | 10.6 | ATCGG | D-loop | n.a. |
| 5 | Asp-GTC | **98.3** | GTCAC | Anticodon loop | 3 |
|  |  | 10.8 | CACGC | Anticodon loop | n.a. |
| 6 | Glu-TTC | **98.5** | TTCAC | Anticodon loop | 3 |
| 7 | Gly-TCC | 26.0-26.4 | TTCCA | Anticodon loop | 2 |
|  |  | 79.4-80.5 | TCCAA | Anticodon loop | 3 |
| 8 | His-GTG | 19.5 | GG**C**CG | Anticodon loop | n.a. |
| 9 | Leu-TAA | 13.3 | TTCAT | Variable loop | n.a. |
| 10 | Leu-TAG | 37.5-73.2 | TTCGG | Variable loop | n.a. |
|  |  | 41.6 | TTCGA |  |  |
| 11 | Lys-CTT | 73.1 | CC**C**TT | Anticodon loop | 1 |
|  |  | 14.5 | CT**C**TT |  |  |
|  |  | 10.0 | CTCGG | D-loop | n.a. |
| 12 | Met-CAT | 24.4-50.8 | CTCAT | Anticodon loop | 1 |
| 13 | Sec-TCA | 46.3 | TTCAA | Anticodon loop | 2 |
|  |  | 84.6 | AG**C**GA | Variable loop | n.a. |
| 14 | Ser-AGA | 13.4 | CT**C**CC | Variable loop | n.a. |
| 15 | Ser-CGA | 12.9-29.5 | CTCGA | Anticodon loop | 1 |
| 16 | Val-CAC | 10.4-11.6 | CT**C**AC | Anticodon loop | 1 |
|  |  | 13.3-18.4 | CACAC | Anticodon loop | 3 |
| 17 | Val-TAC | 47.9-57.7 | TACAC | Anticodon loop | 3 |
|  |  | 11.0 | CACGC | Anticodon loop | n.a. |

(\*) average editing efficiency or a range of average editing efficiency (if edited reads were mapped to more than one isodecoder). The highest editing is marked in bold
(\*\*) deaminated C is marked in bold, (n.a.) not applicable

Taken together, our data indicate that a wide spectrum of tRNA isoacceptors are susceptible to A3A-mediated editing. The latter process preferentially occurs in the anticodon loop and shows high efficiency and site-specificity. The structural requirements of A3A activity on tRNA are in line with those reported for other substrates [14, 21–23].

To verify the above observations indicating site specificity of the A3A-mediated tRNA deamination two synthetic tRNAs (tRNA-Asp-GTC and tRNA-Gly-GCC) were treated with WT_A3A or E72A_A3A. The reaction products were subjected to RT-PCR, cloned, and 20 independent clones from each reaction were sequenced (Figure 5) (for details see Methods). As shown in Figure 5, in tRNA-Asp-GTC there are 20 Cs, 16 of them are located in double-stranded regions and 4 in single-stranded regions. In tRNA-Gly-GCC there are 20 Cs, 15 in double-stranded regions and 5 in single-stranded regions. Our analysis revealed that A3A almost selectively edited Cs located in the third position of the anticodon in both tested tRNAs (C_36_ in tRNA-Asp-GTC and C_35_ in tRNA-Gly-GCC). In addition, in one clone of tRNA-Gly-GCC, C_37_ located in anticodon loop was deaminated. We also observed that the efficiency of editing depended on the sequence context in which the deaminated Cs were located. tRNA-Asp-GTC contained 5’UC_36_ motif which has been reported to be preferred by A3A [22] and tRNA-Gly-GCC contained unfavorable by A3A, 5’CC_35_ motif. Accordingly, C placed in the 5’UC_36_ motif were deaminated in all sequenced clones (editing efficiency was 100%) and in 5’CC_35_ motif were deaminated in 6 out of 20 clones (editing efficiency was 30%). These results indicate that *in vitro*, A3A editing capacity was restricted to the anticodon loop of both synthetic tRNAs.

**Figure 5.**
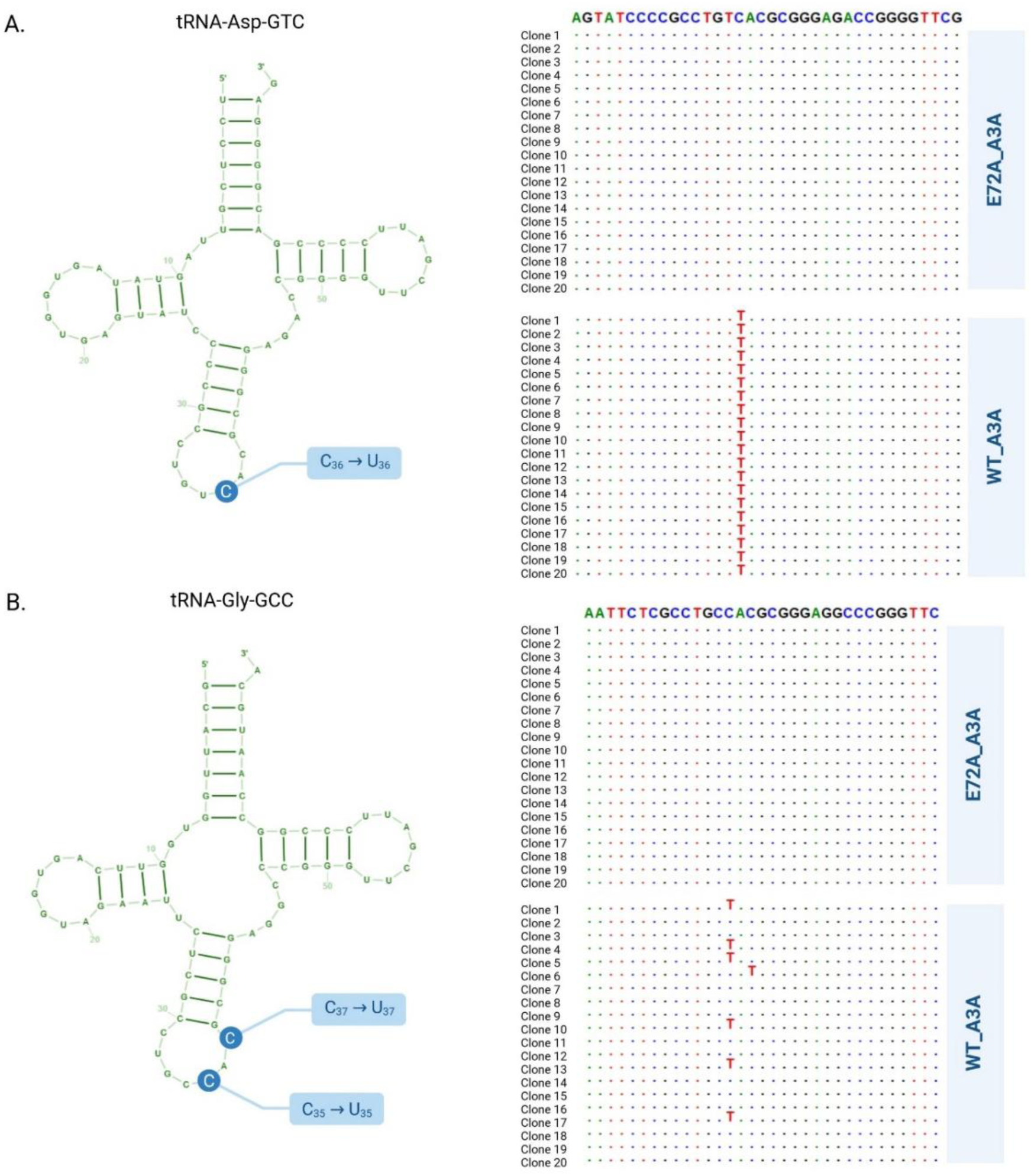
WT_A3A-mediated editing of synthetic tRNAs. Two synthetic tRNAs: tRNA-Asp-GTC (A) and tRNA-Gly-GCC (B) were subjected to editing with WT_A3A. As a control, analogous reactions with E72A_A3A were performed. Reaction products were subjected to RT-PCR, cloned, and 20 randomly selected clones were sequenced by the Sanger method. The locations of edited Cs are schematically indicated in the left panels. The sequencing reads are shown in the right panels (the fragments of tRNA-Asp-GTC sequence located between G20 and A57 and a fragment of tRNA-Gly-GCC sequence located between G21 and G55 are indicated). Unchanged nucleotides are represented by dots. In case of edited nucleotides, blue dots corresponding to C were replaced with T. Created in BioRender. Hoffa-Sobiech, K. (2025) https://BioRender.com/30nsqyc.

### Structural determinants of A3A-mediated tRNA deamination

Intrigued by the ability of A3A to deaminate certain tRNAs, we aimed to elucidate the structural determinants that make particular tRNAs and their anticodon nucleobases optimal or suboptimal substrates. In the absence of high-resolution structures of A3A:tRNA complexes, we employed molecular modeling followed by molecular dynamics (MD) simulations. Three tRNAs (Lys-CTT-3-1, Arg-TCG-1-1, Glu-TTC-3-1) were selected for analysis. The editable C and adjacent tRNA regions were placed in the A3A active site, based on the structure of its complex with ssDNA (PDB ID: 5keg). Initial models confirmed that editing at all three anticodon positions (34-36) was feasible, as no steric clashes were observed between the protein and tRNA (Figure 6A). However, MD simulations revealed variations in the patterns of A3A-tRNA binding (Figure 6B). Glu-TTC-3-1 formed significantly more interactions with A3A than the other two tRNAs (Figure 6B). This appears to result from the initial orientation of the complex, in which A3A was positioned closer to the tRNA core. As a result, the L3 loop (particularly residues Asn61 to Cys64) could directly contact the G10-U11 and A24-C27 regions of the tRNA (Figure 6C). In contrast, for Arg-TCG-1-1, the number of interactions remained comparable to Lys-CTT-3-1, which was edited to a lesser extent. In the Arg-TCG-1-1 model, the L3 loop was positioned close to the anticodon region but did not make as many contacts as in the Glu-TTC-3-1 complex. However, the spatial arrangement suggested that stronger interactions could potentially form if the tRNA molecule were more flexibly bent toward the protein surface. In contrast, in the Lys-CTT-3-1 model, the tRNA remained in a more extended position, and the orientation of A3A limited the loop’s ability to reach the anticodon region, reducing the potential for such interactions.

**Figure 6.**
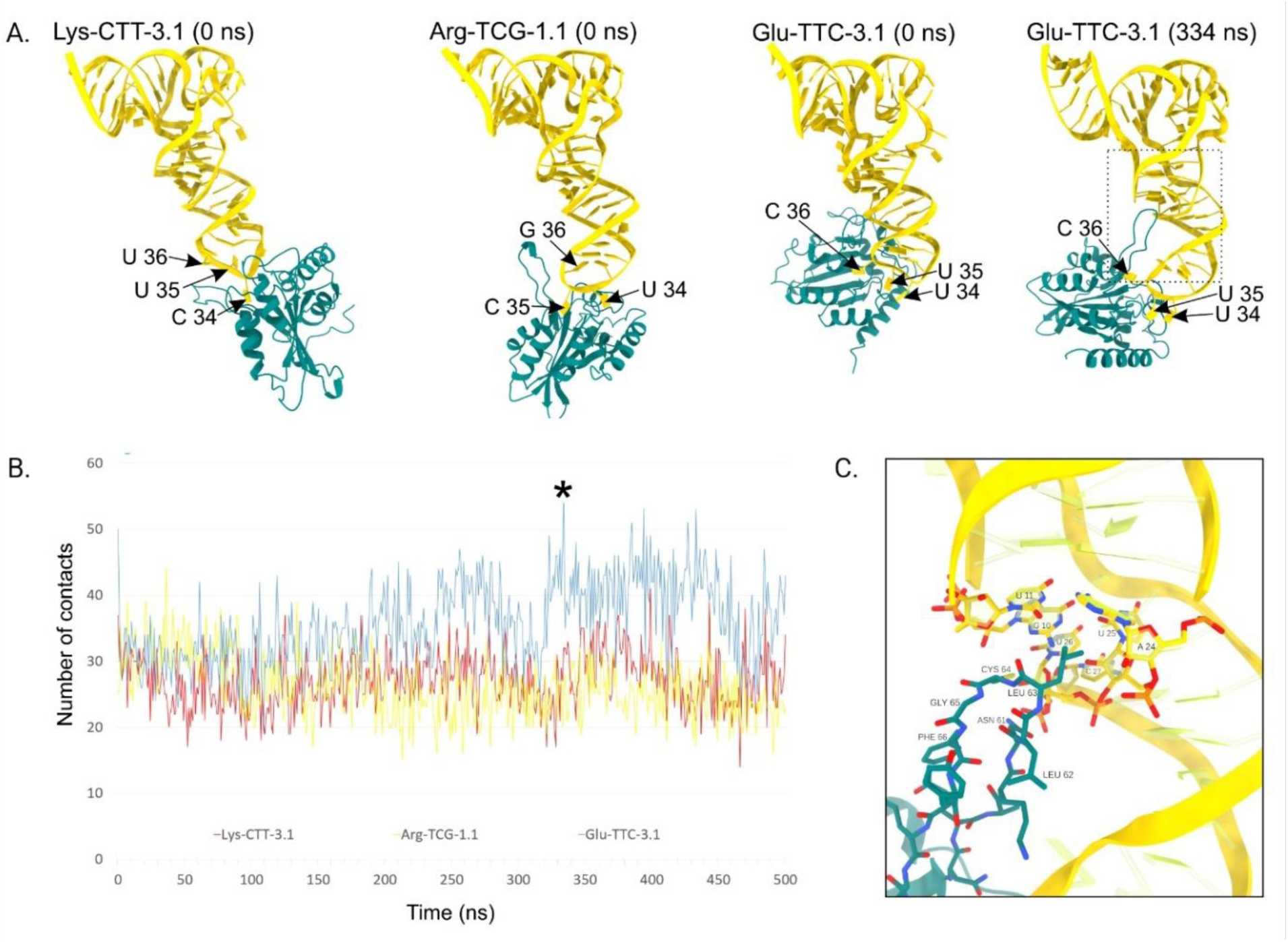
Anticodon nucleobases binding by A3A. (A) Models of A3A structure in a complex with three tRNAs (Lys-CTT-3-1, Arg-TCG-1-1, Glu-TTC-3-1) at the start point of MD simulation (0 ns). Additionally, 334^th^ ns of the A3A:Glu-TTC-3-1 simulation, characterized by the highest number of A3A-tRNA contacts is presented. (B) The number of contacts observed for each A3A:tRNA model throughout MD trajectories. 334^th^ ns of the Glu-TTC-3-1 simulation is marked with an asterisk. (C) Close-up view of the contact region between the L3 loop of A3A and Glu-TTC-3-1 in 334^th^ ns of the simulation. Created in BioRender. Budzko, L. (2025) https://BioRender.com/cbeoirf.

Additionally, we investigated the role of three other loops surrounding the A3A active site—L1, L5, and L7—in forming contacts with tRNA. Among them, the L1 loop showed substantial involvement in all three A3A:tRNA complexes (Supplementary Figure S4), consistent with recent findings on A3A-mediated mRNA recognition [46]. However, unlike the L3 loop, L1 did not show variability in the number of contacts across the models. This suggests that although L1 contributes to A3A:tRNA complex formation in general, L3 loop interactions are likely the key determinant of strong binding at position 36. This supports the hypothesis that L3-mediated interactions may underlie the substrate selectivity of A3A toward specific tRNAs.

### APOBEC-mediated editing is detectable in tRFs in tumor tissues

There are many reports indicating that the *A3A* gene is often upregulated in cancer cells. Moreover, the observation presented above clearly indicates that A3A is capable of editing tRNA. If this process also occurs *in vivo*, then one can expect that A3A-specific editing products should be present in cancer cells. To verify this hypothesis, we searched for these types of products in human cancer samples for which miRNA-Seq, RNA-Seq, and WGS datasets are available in TCGA. Together, we analyzed 1230 samples from 19 cancer types (Supplementary Table S3). Unfortunately, standard approaches used for transcriptome analysis omit tRNA sequencing. However, despite the lack of mature tRNA sequence data, we could detect the tRNA editing products by analyzing tRFs that are sequenced together with the miRNA fraction. To identify editing events in patient-derived samples, we used the list of high-confidence *in vitro* editing sites (Supplementary Table S2) to screen miRNA-seq data from TCGA. These datasets include short RNAs ranging from 15 to 30 nucleotides in length. We excluded any candidate sites that overlapped with somatic or germline variants detected in WGS data. Given the well-known challenges in detecting and mapping tRFs—even in the absence of editing—we further limited our analysis to genomic coordinates corresponding to annotated tRFs in MINTbase v2.0 [47]. As shown in Supplementary Table S4 and Figure 7, we identified six editing positions in the miRNA-seq data. All six sites had sufficient coverage (>10x) in both the miRNA-seq and WGS datasets, were not mutated at the DNA level, and overlapped with annotated tRFs in MINTbase. Three out of six edited sites were located in the second anticodon position, one in the wobble position, one in the third anticodon position, and one in the variable loop. The mutation of tRFs originated from tRNA-Ala-TGC-4-1 (the third anticodon position) and tRNA-Arg-CCG-2-1 (the wobble position) were the most frequent, as they were detected in 151 and 146 patients, respectively. To analyze editing events within individual cancer types, we included only those sites that were detected in at least five patients. This filtering step reduced the number of editing sites under consideration to three (tRNA-Ala-TGC-4-1, tRNA-Arg-CCG-2-1, tRNA-Cys-GCA-5-1). Average editing for these three positions in analyzed patients was 17.8%, 38.5%, and 7.4% for tRFs originating from tRNA-Ala-TGC-4-1, tRNA-Cys-GCA-5-1, and tRNA-Arg-CCG-2-1, respectively (Supplementary Tables S4 and S5). They occurred predominantly in similar cancer types, with the highest number of edited samples detected for THCA (86 samples with cumulative editing for three positions), LUAD (29 samples), HNSC (24 samples), BRCA (15 samples), and OV (15 samples). The average editing was different in specific cancer types, reaching 6-54% for tRFs originating from tRNA-Ala-TGC-4-1, 17-60% for tRFs originating from tRNA-Cys-GCA-5-1, and 2-15% for tRFs originating from tRNA-Arg-CCG-2-1 (Supplementary Table S6).

**Figure 7.**
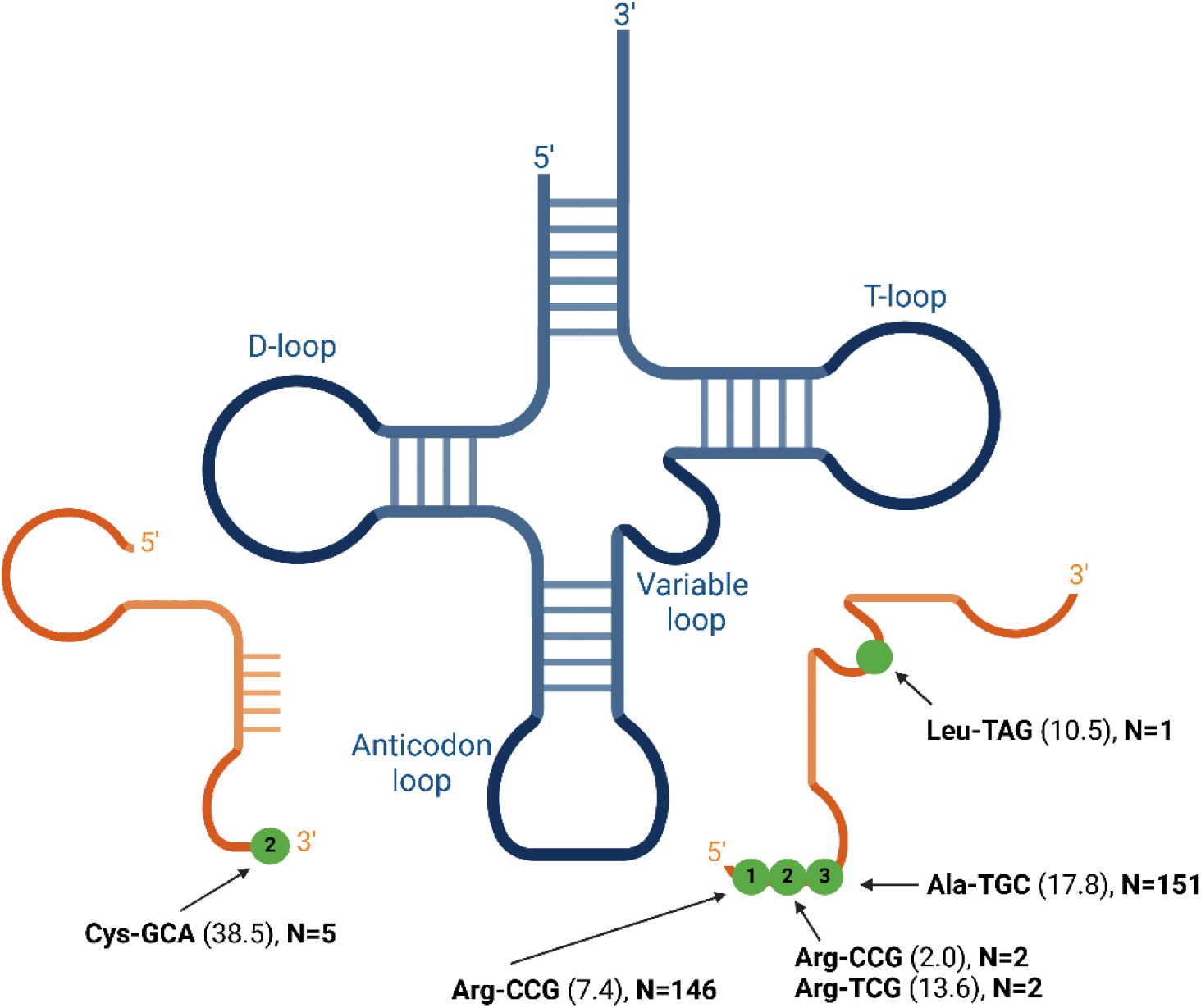
Schematic representation of C>U edited positions identified in tRFs in cancer samples. The green dots represent Cs deaminated in different positions. The average editing efficiency is given in parentheses. N – number of patients found to have the editing in the given position. Created in BioRender. Hoffa-Sobiech, K. (2025) https://BioRender.com/i4pg8hr.

Next, RNA-Seq data were used to analyze *A3A* gene expression level in the samples with detected editing. The *A3A* gene was found to be expressed (> 0.5 Transcripts Per Million) in 66.4% of these samples (denoted A3A+; samples without detectable *A3A* expression were denoted A3A-). As shown in Supplementary Table S6, the detected C>U changes, for all cancers analyzed collectively, were significantly more frequent in A3A+ than in A3A-samples. The analysis was then repeated for specific cancer types, for which sufficient numbers of samples were available. The same dependency was observed for most of the cancer types: BRCA, CESC (cervical squamous cell carcinoma and endocervical adenocarcinoma), HNSC, LUAD, LUSC (lung squamous cell carcinoma), and OV (Supplementary Figure S5 and Supplementary Table S6). Exceptions to this rule were THCA and LGG (brain lower grade glioma). In these cases, many tumor samples with detected editing in tRFs no longer expressed *A3A* at the time of mRNA analysis. However, it should be taken into account that tRFs can be quite stable and persist for long periods, unlike some other types of RNA, such as mRNA [40, 48, 49]. Moreover, a protein showing significant stability can also persist in the cell even when the level of its mRNA has already declined. Finally, we tested whether *A3A* gene expression level correlates with the efficiency of editing detected in tRFS in A3A+ samples. We did not observe such a positive correlation, which suggests that the *A3A* gene expression level cannot predict the editing efficiency in bulk sequencing (Supplementary Figure S6). This result is consistent with recent reports showing that in the human genome, APOBEC-signature mutations are generated in an episodic manner and do not correlate with the expression levels of APOBECs [23, 50]. Taken together, our data suggest that dysregulation of A3A in tumor samples can lead to the accumulation of C>U mutations in tRFs.

## Discussion

To verify our hypothesis that A3A is involved in tRNA editing in humans, we tested A3A capacity to deaminate tRNA *in vitro* and *in vivo*. As a result, we demonstrated that recombinant A3A exhibits C>U deamination activity *in vitro* on synthetic and cell-derived tRNAs. Our results indicated that A3A-mediated tRNA editing is highly effective and site-specific. WT_A3A efficiently (≥10%) deaminated Cs in 17 different tRNA isoacceptors in all three anticodon positions, at the 3’ end of the anticodon loop (position 38), within the variable, and D loops. The third and second anticodon positions were the most frequently (in total in 10 isoacceptors) and the most efficiently edited (max. 98-99% in both cases). Interestingly, their deamination changes the recognized codon sequence (Figure 8). For instance, the identified editing of C_36_>U_36_ in tRNA-Asp-GTC changes the recognized codon sequence from GAC to AAC (Asp- to Asn-coding). Similarly, C_35_>U_35_ editing in tRNA-Arg-CCG alters the recognized codon sequence from CGG to CAG (Arg- to Gln-coding). Therefore, deamination of the second and third anticodon positions (Figure 8A, B) or both at the same time (Figure 8C) potentially contributes to the mistranslation. However, different effects can be expected in the case of deamination of aminoacyl-tRNA (aa-tRNA or charged tRNA) and non-charged tRNA. Although *in vivo* A3A specificity towards charged- and non-charged tRNA remains elusive, one can hypothesize the following scenarios. The aa-tRNA deamination may lead to the recognition of a different codon and incorporation of an amino acid not specified by the genetic code. On the other hand, deamination of non-charged tRNA may disrupt the activity of aminoacyl-tRNA synthetases (AARS). Mutations within the anticodon (which is one of the identity elements of tRNA) are likely to either inhibit AARS function or cause the tRNA to acquire a new identity, leading to incorrect aminoacylation. For instance, it has been shown that a substitution of one C in the variable loop of *S. cerevisiae* tRNA-Leu-TAG destabilizes the tetra-loop structure and leads to the incorrect charging of this tRNA with Ser [51]. The effect of substitution is even more pronounced when it occurs in the anticodon. It has been demonstrated that the correct aminoacylation of tRNA-Met, tRNA-Arg, tRNA-Glu, and tRNA-Ile requires specific recognition of the anticodon nucleobases by AARS, and even a single-nucleotide change within the anticodon can disrupt these interactions [52–56].

**Figure 8.**
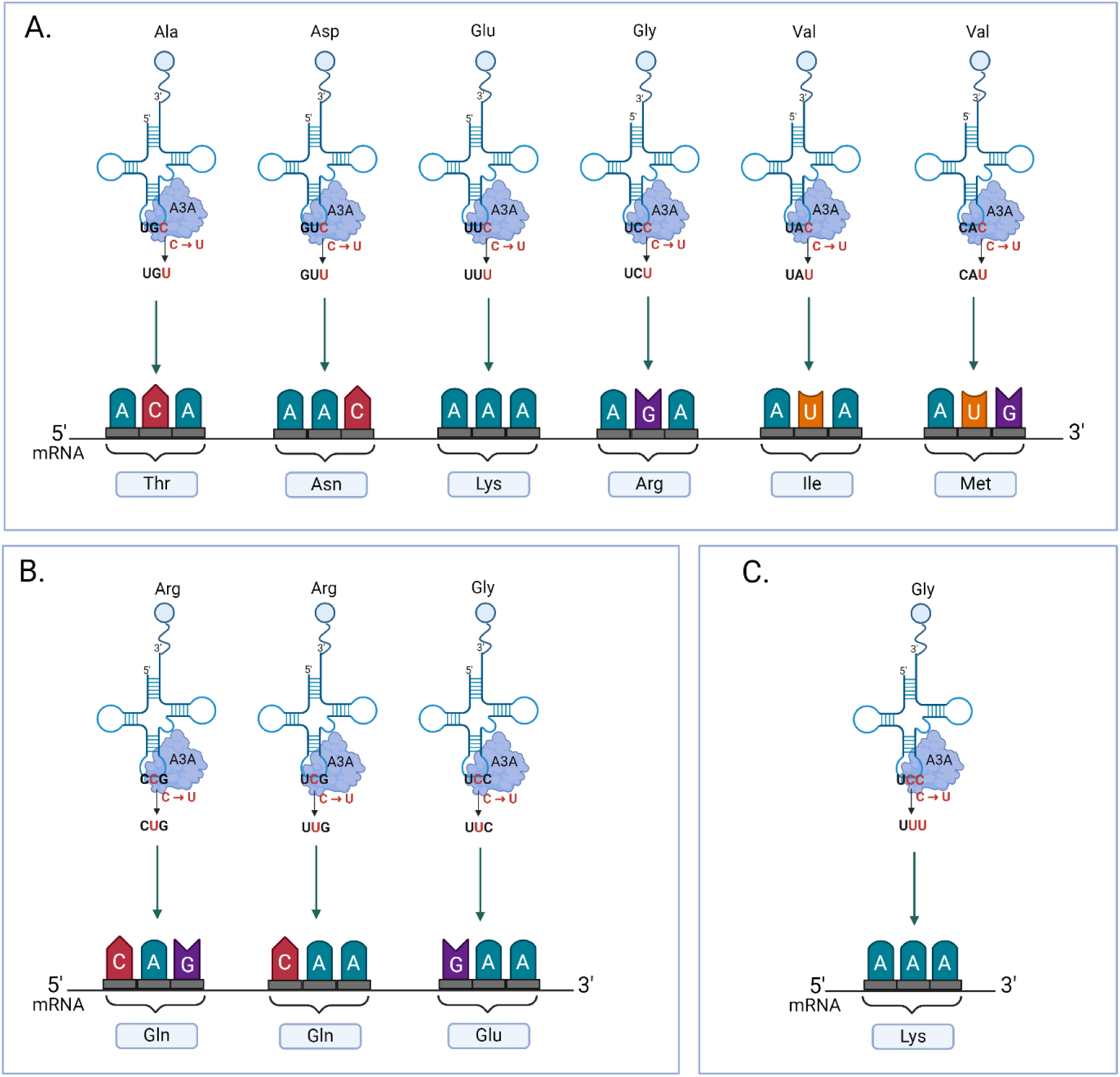
Changes in the recognized codon sequence resulting from the A3A-mediated deamination of the third (A) and second (B) anticodon position, or both (C). Created in BioRender. Hoffa-sobiech, K. (2025) https://BioRender.com/eg6ezzs.

The editing of the first anticodon position occurred less frequently (in four isoacceptors) and was less efficient in our study (max. 73%). Since the wobble position exhibits flexibility in base pairing (i.e. it only distinguishes between purine or pyrimidine in the codon), its deamination generally does not change the recognized codon sequence. An exception is tRNA-Lys-CTT whose deamination observed in our data may result in the change of the recognized codon sequence from AAG to AAC (from Lys- to Asn-coding). However, deamination of the wobble position may have potentially other important consequences. For instance, it may influence the biogenesis and functions of post-transcriptional modifications, abundantly occurring at this position [57]. One can speculate that C>U deamination may either promote a biogenesis of U-modifications (e.g. pseudouridine (Ψ) formation [34]) or disturb the pathways of C-modification (e.g. methylation of C to 5-methylcytidine (m5C) [58]). The modifications of wobble position are implicated in maintaining accurate decoding, translational processivity, and preventing frameshifting [57, 59, 60]. Therefore, the deficiencies in these modifications can have potentially pathological consequences, such as the reported: mitochondrial diseases, neurological disorders, and cancer [59, 61, 62]. Noteworthy, viral RNA sequences often differ significantly from host cell mRNAs, with many viruses favoring A/U-ending codons over the C/G-ending codons prevalent in the host genome [63, 64]. Therefore, the wobble position editing may increase the pool of tRNA species capable of decoding A-ending codons.

For several tRNA isoacceptors, we observed A3A-mediated deamination outside the anticodon. For three tRNA isoacceptors, we observed deamination at position 38, two tRNA isoacceptors were deaminated within the D-loop (position 17), and four within the variable loop. Importantly, some of these positions are known for being post-transcriptionally modified. More specifically, Ψ and m5C are characteristic of position 38 in the anticodon loop, and dihydrouridine (D) is characteristic of position 17 of the D-loop [65–67]. Although research on modifications in the variable loop is relatively sparse compared to other tRNA regions, there are indications of possible pseudouridylation and C-methylation of these positions [68, 69]. Similarly, as in the case of the anticodon deamination, editing of the position 38, D-loop, or variable loop may have a significant impact on tRNA-processing (e.g. by AARS, modifying enzymes, or ribonucleases) further affecting translation effectivity and fidelity.

The observed in our data characteristics of A3A activity on tRNA were consistent with those previously reported for A3A activity on other substrates [14, 22, 46]. In both synthetic and cell-derived tRNAs deamination occurred within lops of stem-loop structures and sites undergoing A3A-mediated tRNA editing showed an enrichment of the 5΄YU<u>C</u>R motif. However, the example of tRNA-Sec-TCA, in which an unfavorable sequence context within the variable loop was more effectively edited than a favorable context within the anticodon, suggests that other structural factors may also play a crucial role in determining the editing efficiency. Therefore, to identify the mechanism by which A3A recognizes specific anticodon positions in different tRNAs, we employed molecular modeling followed by MD simulations of A3A bound to different anticodon positions in three tRNAs (Lys-CTT-3-1, Arg-TCG-1-1, Glu-TTC-3-1). Our analysis revealed the highest number of A3A:tRNA contacts for Glu-TTC-3-1 having C in the third anticodon position. The largest contribution to the formation of A3A:tRNA contacts was made by the L1 and L3 loops. However, the L3 loop was the differentiating factor in the number of contacts between the three generated models. Notably, the L1 loop has been recently proposed to be an important factor in mRNA recognition by A3A [46], and other studies confirm the involvement of the L1, L3, and L7 loop in DNA substrate recognition [5, 6, 70]. Future structural studies on tRNA, preferentially in the complex with A3A, will likely deepen our understanding of A3A:tRNA recognition.

Through bioinformatic analysis of sequencing data deposited in TCGA, we showed that A3A-mediated tRNA mutations observed *in vitro* can be found in tRFs accumulated in the cancer tissues. Comparing miRNA-Seq, RNA-Seq, and WGS data, we identified six residues edited by A3A *in vitro* which were also present in cancer-accumulated tRFs but were not mutated in the corresponding WGS data. Notably, three of them were found in a significant number of patient samples. Although the A3A-mediated tRNA/tRF editing in cancer cell lines remains elusive, it can be speculated that deamination may also impact tRF biogenesis and/or function. Interestingly, two main tRF classes - tRNA halves (5’-halves and 3’-halves) are generated by the cleavage at the anticodon of mature tRNAs by angiogenin (ANG) [71–75]. Therefore one can speculate that C>U deamination within the anticodon nucleobases may influence their recognition and cleavage by ANG. Moreover, the change of tRF sequences may influence their regulatory functions such as the well-documented miRNA-like function [76–79].

Mistranslation has been described as a result of impaired expression of genetic information at various stages [80]. This phenomenon has been well documented during viral infections and cancer development [81, 82], and *A3A* is upregulated in both these pathological conditions [83, 84]. It has been hypothesized that viruses utilize mistranslation for viral evolution [85]. Mistranslation can also result in the production of host proteins with altered structure and function, potentially disrupting key cellular processes and promoting cancer development [86–88]. Previously, it has been shown that A3A modifies genetic information by introducing mutations to the human genome and mRNAs [16, 23]. This study shows that A3A may interfere with the flow of genetic information also by tRNA editing, which sheds new light on A3A’s role in both viral infection and oncogenesis. Importantly, apart from being an aberrant process, mistranslation is actively induced under certain circumstances, providing adaptive advantages by generating protein diversity and allowing cells to rapidly respond to environmental changes [89–91]. Therefore, tRNA deamination may also have yet unknown beneficial consequences, the specifics of which remain to be elucidated.

In conclusion, our finding that A3A deaminates tRNA *in vitro* opens new avenues of research on RNA editing phenomenon and mistranslation in the context of both physiological and pathological processes. We showed for the first time that A3A efficiently deaminates a large spectrum of tRNA isoacceptors and shows high specificity towards anticodon nucleobases. Moreover, our bioinformatic analysis confirmed that we can find traces of this phenomenon in sequencing data from human cancer tissues in which A3A is deregulated. Although the A3A-mediated tRNA editing *in cellulo* remains elusive, our data suggest that A3A activity may disturb the decoding capacity of a large pool of tRNAs, as well as influence potentially all processes related to tRNA functions. Our finding has also far-reaching implications for discovering new tumor markers and the development of antiviral therapies.

## Materials and methods

### Cloning, expression, and purification of WT APOBEC3A and its inactive mutant

The genes encoding human WT_A3A (Genbank No NM_145699.4) and its catalytically inactive mutant - E72A_A3A generated by site-directed mutagenesis were cloned into pET42A(+) (Novagen) and pGEX4T1 (Novagen) expression vectors, respectively. Both expression constructs contained sequences encoding: GST, a thrombin cleavage site, His-tag, and an enterokinase cleavage site, in that order. The sequences of all primers are available upon request. Both constructs were transformed into BL21(DE3)pLysS cells (Novagen). WT_A3A protein production was conducted for 2 h at 37 °C, while E72A_A3A production was carried for 16 h at 18 °C, after induction of overexpression with isopropyl β-D-1-thiogalactopyranoside (IPTG) in the final concentration of 0.2 mM and 0.5 mM, respectively. Additionally, growth media were supplemented with 0.06 mM zinc chloride. The bacterial culture was centrifuged at 3500xg for 20 min at 4 °C and stored at −80°C for purification. WT_A3A and E72A_A3A were pre-purified using Glutathione Sepharose 4 Fast Flow column (GE Healthcare) according to the manufacturer’s protocol, followed by overnight cleavage with thrombin protease (Merck) at 4 °C with simultaneous dialysis into the filtration buffer (50 mM Hepes, pH=7.2, 1mM TCEP, 0.1M NaCl). Native enzymes (stripped of tags) were purified by size-exclusion chromatography performed using Superdex 75 16/60 column (GE Healthcare) and the ACTA pureTM chromatography system (Cytiva). Fractions eluted with a peak at 23 kDa were concentrated, analyzed on SDS-PAGE (Supplementary Figure S1), and used in the deamination assays. All efforts were made to prevent contamination of the WT_A3A enzyme in the preparation of the mutant and vice versa (for example, different columns for purification were used).

### Deamination of DNA and RNA oligonucleotides

All sequences of DNA and RNA oligonucleotides used in the deamination reaction are listed in Supplemental Table S7. For the deamination assay, 3 μg of each DNA or RNA oligomer was denatured (followed by refolding by slow cooling down to 4 °C in the case of RNA oligomers) and incubated for 3 h at 37 °C with 4 μg of WT_A3A or E72A_A3A mutant, in the reaction buffer (50 mM MES, pH=6,0, 1 mM TCEP, 0,1M NaCl). After reaction, oligonucleotides were digested to nucleosides using Nucleoside Digestion Mix (New England Biolabs) according to the manufacturer’s protocol and analyzed with UPLC.

### Nucleoside analysis with ultra performance liquid chromatography (UPLC)

DNA and RNA hydrolysates were mixed with 5uL of MilliQ-grade deionized water and ultrafiltered prior to injection using filter plates with modified polyethersulfone membranes (10K). DNA hydrolysates were spiked with internal standards in a volumetric ratio 4:1, to a concentration of 1 pmol/µL of [^13^C_9_,^15^N_3_]-dC and 50 fmol/µL of [^13^C,^15^N_2_]-dU, and analyzed using 2D-UPLC-MS/MS. Chromatographic separation was achieved using a method described by Starczak et al. [92] based on a 2D-UPLC system with photo-diode array detector for the first dimension chromatography (used for the quantification of 2’-deoxythymidine (dT) and 2’-deoxyguanosine(dG)) and tandem quadrupole mass spectrometer (Xevo TQ-XS, Waters) using the following columns: Waters Cortecs T3 column (150 mm×3 mm, 1.6 µm) with a precolumn at the first dimension, a Waters X-select C18 CSH (100 mm×2.1 mm, 1.7 µm) at the second dimension and Waters X-select C18 CSH (20 mm×3 mm, 3.5 µm) as a trap/transfer column. At the first dimension a flow rate of 0.5 mL/min was used, with an injection volume of 2 µL and gradient elution for 10 min using a mobile phase of 0.05% acetate (A) and acetonitrile (B) (0.7-5% B for 5 min, followed by the column washing with 30% acetonitrile and re-equilibration with 99% A for 3.6 min). At the second dimension the flow rate was 0.35 mL/min in a gradient elution for 10 min using a mobile phase of 0.01% acetate (A) and methanol (B) (1-50% B for 4 min, isocratic flow of 50% B for 1.5 min, and re-equilibration with 99% A up to the next injection). The quantities of dT and dG were determined by UV detection at 260 nm and 280 nm, respectively.

Chromatographic separation of RNA samples was performed using a UPLC system with a photo-diode array detector for the quantification of canonical ribonucleosides, using Waters Cortecs T3 column (150 mm×3 mm, 1.6 µm) with a precolumn. Flow rate was 0.5 mL/min, with an injection volume 2 µL and gradient elution was applied for 10 min using a mobile phase 0.05% acetate with 5µM ammonium formate (A) and acetonitrile (B) (0.7% B for 0.5 min, followed by gradient (0.7-50% B) for 4.5 min, the column was washed with 50% acetonitrile for 2.2 min and re-equilibrated with 99% A for 2.8 min. The quantities of canonical ribonucleosides were determined by UV detection at 260 nm for rU, 280 nm for guanosine, adenosine, and rC. The UV peaks for rC and rU peaks were confirmed by monitoring of MS traces: 244→112 (rC) and 243→110 (rU).

### HEK293T cell culture and RNA isolation

Human embryonic kidney cells (HEK293T) were cultured as described in Supplementary Methods. Total RNA was extracted from snap-frozen cells pellet of HEK293T using mirVana^TM^ miRNA isolation kit (Invitrogen) according to the manufacturer’s protocol. The integrity of total RNA was assessed using the Agilent TapeStation System. Samples were enriched in small RNA fraction (<200 nt) using mirVana^TM^ miRNA isolation kit. For purification of tRNA fraction small RNA was electrophoresed on 8% Urea-TBE denaturing polyacrylamide gel. The target fraction of tRNA was visualized using SYBR-Gold (ThermoFisher Scientific) and excised (70 – 90 nucleotides in length) under UV light. The excised tRNAs were extracted using sodium acetate and precipitated. tRNA purity and integrity were confirmed using a NanoDrop1000 spectrophotometer and Agilent TapeStation System.

### Sanger sequencing analysis of deaminated synthetic tRNAs

Sequences of tRNA oligonucleotides (tRNA-ASP-GTC-2-1, tRNA-Gly-GCC-2-1; synthesized by Future Synthesis), used in the deamination reaction, are listed in Supplemental Table S7. Synthetic tRNAs were denatured in the reaction buffer (50 mM MES, pH=6.0, 1 mM TCEP, 0.1M NaCl, 5 mM MgCl_2_) for 4 minutes at 95 °C, and slowly cooled down to 4 °C. For the deamination assay, re-folded tRNAs were incubated for 3 h at 37 °C with 4 μg of WT_A3A or E72A_A3A mutant, in the reaction buffer. Products of the deamination reaction were purified using MinElute PCR Purification kit (Qiagen), followed by one-step reverse transcription-polymerase chain reaction (RT-PCR) using SuperScript IV One-Step RT-PCR System (ThermoFisher Scientific) according to the manufacturer’s protocol. RT-PCR products were cloned into pCR Blunt II-TOPO vector (ThermoFisher Scientific). Twenty individual randomly selected clones were sequenced per reaction. Subsequently, the obtained sequences were aligned and compared with the sequence of the original oligonucleotide.

### Deamination of mature tRNA and preparation of libraries for NGS sequencing

The fraction of mature tRNA was isolated from HEK293T cells as described above. tRNA fraction was deaminated (as described above for synthetic tRNAs), followed by purification using the Oligo Clean & Concentrator kit (Zymo Research). Subsequently, tRNA was incubated at 37°C for 40 min in 0.1 M Tris-HCl (pH=9) to remove amino acids (deacylation treatment), followed by ethanol precipitation. tRNA libraries were prepared according to the previously described YAMAT-seq protocol [45]. The quality of libraries was confirmed using the Agilent TapeStation System. Libraries were sequenced on NovaSeq 6000 Illumina platform.

### Computational analysis of tRNA editing in NGS sequencing data

The numbers of raw reads obtained from libraries prepared according to the YAMAT-seq protocol for WT_A3A- or E72A-treated samples and untreated (with technical triplicates) are shown in Supplementary Table S8. A detailed description of the computational analysis is available in Supplementary Methods. Briefly, after trimming, filtering, and quality check, the libraries were normalized by the number of reads and 40 mln reads for each library were randomly selected, and a selected read could be chosen only once. The reads were then aligned to the hg38 human genome. We allowed non-unique mappings of reads and filtered out reads with more than 3 mismatches. Reads were also excluded if MAPQ was lower than 20. During the mapping we used alignments scoring and chose the match with the highest score. Next, the aligned reads that exceeded the tRNA boundary by at least 1 bp were removed (tRNA-reference genes were listed in gtRNAdb [93] release 21). We considered a tRNA being present in the sample if the minimum coverage of its genomic coordinates was 250×. Next, variant calling was performed on strand-specific BAM files. Sets of C>T (for positive DNA strand) and G>A (for negative DNA strand) substitutions were annotated by mapping reads to tRNA genomic regions. To evaluate a position we required at least 1% of reads supporting an alternate allele (minimum 3 reads). A position was considered to be edited if the mismatch was detected for at least two out of three replicates among the WT_A3A-treated samples. These sites were processed to calculate editing efficiency and the fold change of detected editing between pairs of WT_A3A-treated and control samples (separately for E72A control and untreated). WT_A3A-modified sites in comparison to both controls whose fold-change was equal or higher than 2 were considered to be edited positions and used for bootstrap analysis. The bootstrap analysis with ∼1000 resamples was performed to investigate the replicating probability of obtained results. The editing site was considered as high-confident if: (i) the resampling had a minimum of 95% bootstrap accuracy; (ii) the mean editing of the primary call was between the lower and upper bound of bootstrap distribution with 95% CI.

### Molecular modeling of A3A:tRNA complex and molecular dynamics simulations

The models of A3A complexes with three representative tRNAs (Lys-CTT-3.1; Arg-TCG-1.1; Glu-TTC-3.1) were built using the following protocol. Unbound tRNAs were modelled using the Alphafold 3 server [94]. Then, the position and conformation of the region of interest with respect to A3A were modeled based on the PDB ID: 5keg entry; the modified C was placed in the catalytic site. Such models were imported into *Maestro 2024-2* (Schrodinger) and processed using the protein preparation tool, including the addition of hydrogen atoms, optimization of their geometry, and energy minimization.

The MD simulations were performed in *Desmond 7.8.134* [95], within *Maestro* 2024-2. Each system was configured with the OPLS4 force field [96] using TIP4P solvent model. The charges were neutralized by adding Na^+^/Cl^−^ ions, and the equivalent of 150 mM NaCl was introduced to the system. A standard relaxation protocol was applied prior to the final simulation runs. Production MD simulations were performed for 500 ns at 300 K in the NPT ensemble, with data recorded every 100 ps, yielding 5,000 frames. Each system was simulated at least twice, using randomized velocities. Every-1-ns snapshots were exported as .pdb files and analyzed using Contact within the CCP4-9 [97] in batches using custom-made scripts. Figures were created in UCSF ChimeraX [98].

### Computational analysis of editing in TCGA data

To identify putative A3A-deaminated positions in miRNA-Seq data deposited in TCGA database (release 36), we analyzed samples for which data sets consisting of miRNA-Seq, RNA-Seq, and WGS data (paired tumor-normal data) were available. In total, we analyzed 1230 samples from 19 cancer types (see Supplementary Table S3). The RNA-seq data were downloaded as the gene expression quantification data type to estimate the A3A expression level. The tumor-normal WGS data were processed to call somatic and germline variants. All data were mapped to the human genome build hg38. The aligned miRNA-Seq reads that exceeded tRNA boundaries by at least 1 bp were removed. The list of *in vitro* editing sites shown in Supplementary Table S2 was used to search the miRNA-seq data. The variant found in TCGA data was selected if the minimum depth for this variant coordinate was 10 in both miRNA-Seq data and WGS tumor-normal datasets. Finally, positions overlapping with somatic and/or germline variants in at least one patient were excluded. A detailed description of the computational analysis is available in Supplementary Methods.

## Acknowledgments

This work was funded by the National Center for Research and Development under the LIDER IX Program (project no. LIDER/30/0111/L9/17/NCBR/2018).

## Reuse note

Reuse of this work is subject to the CC BY-NC-ND license.

## References

1. Conticello, S.G., The AID/APOBEC family of nucleic acid mutators. Genome Biol, 2008. 9(6): p. 229.

2. Salter, J.D., R.P. Bennett, and H.C. Smith, The APOBEC Protein Family: United by Structure, Divergent in Function. Trends Biochem Sci, 2016. 41(7): p. 578–594.

3. Harris, R.S. and M.T. Liddament, Retroviral restriction by APOBEC proteins. Nat Rev Immunol, 2004. 4(11): p. 868–77.

4. Silvas, T.V. and C.A. Schiffer, APOBEC3s: DNA-editing human cytidine deaminases. Protein Sci, 2019. 28(9): p. 1552–1566.

5. Hou, S., et al., Structural basis of substrate specificity in human cytidine deaminase family APOBEC3s. J Biol Chem, 2021. 297(2): p. 100909.

6. Salter, J.D. and H.C. Smith, Modeling the Embrace of a Mutator: APOBEC Selection of Nucleic Acid Ligands. Trends Biochem Sci, 2018. 43(8): p. 606–622.

7. Byeon, I.J., et al., NMR structure of human restriction factor APOBEC3A reveals substrate binding and enzyme specificity. Nat Commun, 2013. 4: p. 1890.

8. Narvaiza, I., et al., Deaminase-independent inhibition of parvoviruses by the APOBEC3A cytidine deaminase. PLoS Pathog, 2009. 5(5): p. e1000439.

9. Warren, C.J., et al., APOBEC3A functions as a restriction factor of human papillomavirus. J Virol, 2015. 89(1): p. 688–702.

10. Ooms, M., et al., APOBEC3A, APOBEC3B, and APOBEC3H haplotype 2 restrict human T-lymphotropic virus type 1. J Virol, 2012. 86(11): p. 6097–108.

11. Ito, F., et al., Family-Wide Comparative Analysis of Cytidine and Methylcytidine Deamination by Eleven Human APOBEC Proteins. J Mol Biol, 2017. 429(12): p. 1787–1799.

12. Langenbucher, A., et al., An extended APOBEC3A mutation signature in cancer. Nat Commun, 2021. 12(1): p. 1602.

13. Silvas, T.V., et al., Substrate sequence selectivity of APOBEC3A implicates intra-DNA interactions. Sci Rep, 2018. 8(1): p. 7511.

14. Buisson, R., et al., Passenger hotspot mutations in cancer driven by APOBEC3A and mesoscale genomic features. Science, 2019. 364(6447).

15. Law, E.K., et al., APOBEC3A catalyzes mutation and drives carcinogenesis in vivo. J Exp Med, 2020. 217(12).

16. DeWeerd, R.A., et al., Prospectively defined patterns of APOBEC3A mutagenesis are prevalent in human cancers. Cell Rep, 2022. 38(12): p. 110555.

17. Sanchez, A., et al., Mesoscale DNA features impact APOBEC3A and APOBEC3B deaminase activity and shape tumor mutational landscapes. Nat Commun, 2024. 15(1): p. 2370.

18. Butler, K. and A.R. Banday, APOBEC3-mediated mutagenesis in cancer: causes, clinical significance and therapeutic potential. J Hematol Oncol, 2023. 16(1): p. 31.

19. Isozaki, H., et al., Therapy-induced APOBEC3A drives evolution of persistent cancer cells. Nature, 2023. 620(7973): p. 393–401.

20. Roberts, S.A., et al., An APOBEC cytidine deaminase mutagenesis pattern is widespread in human cancers. Nat Genet, 2013. 45(9): p. 970–6.

21. Sharma, S., et al., APOBEC3A cytidine deaminase induces RNA editing in monocytes and macrophages. Nat Commun, 2015. 6: p. 6881.

22. Sharma, S. and B.E. Baysal, Stem-loop structure preference for site-specific RNA editing by APOBEC3A and APOBEC3G. PeerJ, 2017. 5: p. e4136.

23. Jalili, P., et al., Quantification of ongoing APOBEC3A activity in tumor cells by monitoring RNA editing at hotspots. Nat Commun, 2020. 11(1): p. 2971.

24. Sharma, S., et al., Transient overexpression of exogenous APOBEC3A causes C-to-U RNA editing of thousands of genes. RNA Biol, 2017. 14(5): p. 603–610.

25. Green, A.M., et al., Interaction with the CCT chaperonin complex limits APOBEC3A cytidine deaminase cytotoxicity. EMBO Rep, 2021. 22(9): p. e52145.

26. Dixit, S., J.C. Henderson, and J.D. Alfonzo, Multi-Substrate Specificity and the Evolutionary Basis for Interdependence in tRNA Editing and Methylation Enzymes. Front Genet, 2019. 10: p. 104.

27. Torres, A.G., et al., Human tRNAs with inosine 34 are essential to efficiently translate eukarya-specific low-complexity proteins. Nucleic Acids Res, 2021. 49(12): p. 7011–7034.

28. Randau, L., et al., A cytidine deaminase edits C to U in transfer RNAs in Archaea. Science, 2009. 324(5927): p. 657–9.

29. Alfonzo, J.D., et al., C to U editing of the anticodon of imported mitochondrial tRNA(Trp) allows decoding of the UGA stop codon in Leishmania tarentolae. EMBO J, 1999. 18(24): p. 7056–62.

30. Janke, A. and S. Paabo, Editing of a tRNA anticodon in marsupial mitochondria changes its codon recognition. Nucleic Acids Res, 1993. 21(7): p. 1523–5.

31. Zhou, W., D. Karcher, and R. Bock, Identification of enzymes for adenosine-to-inosine editing and discovery of cytidine-to-uridine editing in nucleus-encoded transfer RNAs of Arabidopsis. Plant Physiol, 2014. 166(4): p. 1985–97.

32. Charriere, F., et al., Dual targeting of a single tRNA(Trp) requires two different tryptophanyl-tRNA synthetases in Trypanosoma brucei. Proc Natl Acad Sci U S A, 2006. 103(18): p. 6847–52.

33. Rubio, M.A., et al., An adenosine-to-inosine tRNA-editing enzyme that can perform C-to-U deamination of DNA. Proc Natl Acad Sci U S A, 2007. 104(19): p. 7821–6.

34. Kimura, S., et al., Sequential action of a tRNA base editor in conversion of cytidine to pseudouridine. Nat Commun, 2022. 13(1): p. 5994.

35. Marechal-Drouard, L., et al., A single editing event is a prerequisite for efficient processing of potato mitochondrial phenylalanine tRNA. Mol Cell Biol, 1996. 16(7): p. 3504–10.

36. Binder, S., A. Marchfelder, and A. Brennicke, RNA editing of tRNA(Phe) and tRNA(Cys) in mitochondria of Oenothera berteriana is initiated in precursor molecules. Mol Gen Genet, 1994. 244(1): p. 67–74.

37. Marechal-Drouard, L., et al., RNA editing of larch mitochondrial tRNA(His) precursors is a prerequisite for processing. Nucleic Acids Res, 1996. 24(16): p. 3229–34.

38. Rubio, M.A., et al., Editing and methylation at a single site by functionally interdependent activities. Nature, 2017. 542(7642): p. 494–497.

39. Liu, B., et al., Deciphering the tRNA-derived small RNAs: origin, development, and future. Cell Death Dis, 2021. 13(1): p. 24.

40. Fu, M., et al., Emerging roles of tRNA-derived fragments in cancer. Mol Cancer, 2023. 22(1): p. 30.

41. Budzko, L., et al., Engineered deaminases as a key component of DNA and RNA editing tools. Mol Ther Nucleic Acids, 2023. 34: p. 102062.

42. Bohn, M.F., et al., The ssDNA Mutator APOBEC3A Is Regulated by Cooperative Dimerization. Structure, 2015. 23(5): p. 903–911.

43. Logue, E.C., et al., A DNA sequence recognition loop on APOBEC3A controls substrate specificity. PLoS One, 2014. 9(5): p. e97062.

44. Alqassim, E.Y., et al., RNA editing enzyme APOBEC3A promotes pro-inflammatory M1 macrophage polarization. Commun Biol, 2021. 4(1): p. 102.

45. Shigematsu, M., et al., YAMAT-seq: an efficient method for high-throughput sequencing of mature transfer RNAs. Nucleic Acids Res, 2017. 45(9): p. e70.

46. Kim, K., et al., Unraveling the Enzyme-Substrate Properties for APOBEC3A-Mediated RNA Editing. J Mol Biol, 2023. 435(17): p. 168198.

47. Pliatsika, V., et al., MINTbase v2.0: a comprehensive database for tRNA-derived fragments that includes nuclear and mitochondrial fragments from all The Cancer Genome Atlas projects. Nucleic Acids Res, 2018. 46(D1): p. D152–D159.

48. Costa, B., et al., Nicked tRNAs are stable reservoirs of tRNA halves in cells and biofluids. Proc Natl Acad Sci U S A, 2023. 120(4): p. e2216330120.

49. Gambaro, F., et al., Stable tRNA halves can be sorted into extracellular vesicles and delivered to recipient cells in a concentration-dependent manner. RNA Biol, 2020. 17(8): p. 1168–1182.

50. Petljak, M., et al., Characterizing Mutational Signatures in Human Cancer Cell Lines Reveals Episodic APOBEC Mutagenesis. Cell, 2019. 176(6): p. 1282–1294 e20.

51. Himeno, H., et al., Only one nucleotide insertion to the long variable arm confers an efficient serine acceptor activity upon Saccharomyces cerevisiae tRNA(Leu) in vitro. J Mol Biol, 1997. 268(4): p. 704–11.

52. Schulman, L.H. and H. Pelka, Anticodon loop size and sequence requirements for recognition of formylmethionine tRNA by methionyl-tRNA synthetase. Proc Natl Acad Sci U S A, 1983. 80(22): p. 6755–9.

53. Schulman, L.H. and H. Pelka, Anticodon switching changes the identity of methionine and valine transfer RNAs. Science, 1988. 242(4879): p. 765–8.

54. Schulman, L.H. and H. Pelka, The anticodon contains a major element of the identity of arginine transfer RNAs. Science, 1989. 246(4937): p. 1595–7.

55. Jahn, M., M.J. Rogers, and D. Soll, Anticodon and acceptor stem nucleotides in tRNA(Gln) are major recognition elements for E. coli glutaminyl-tRNA synthetase. Nature, 1991. 352(6332): p. 258–60.

56. Pallanck, L. and L.H. Schulman, Anticodon-dependent aminoacylation of a noncognate tRNA with isoleucine, valine, and phenylalanine in vivo. Proc Natl Acad Sci U S A, 1991. 88(9): p. 3872–6.

57. Ranjan, N. and M.V. Rodnina, tRNA wobble modifications and protein homeostasis. Translation (Austin), 2016. 4(1): p. e1143076.

58. Van Haute, L., et al., NSUN2 introduces 5-methylcytosines in mammalian mitochondrial tRNAs. Nucleic Acids Res, 2019. 47(16): p. 8720–8733.

59. Cui, W., et al., tRNA Modifications and Modifying Enzymes in Disease, the Potential Therapeutic Targets. Int J Biol Sci, 2023. 19(4): p. 1146–1162.

60. Yarian, C., et al., Accurate translation of the genetic code depends on tRNA modified nucleosides. J Biol Chem, 2002. 277(19): p. 16391–5.

61. Bento-Abreu, A., et al., Elongator subunit 3 (ELP3) modifies ALS through tRNA modification. Hum Mol Genet, 2018. 27(7): p. 1276–1289.

62. Kirino, Y., et al., Codon-specific translational defect caused by a wobble modification deficiency in mutant tRNA from a human mitochondrial disease. Proc Natl Acad Sci U S A, 2004. 101(42): p. 15070–5.

63. Ahn, I. and H.S. Son, Evolutionary analysis of human-origin influenza A virus (H3N2) genes associated with the codon usage patterns since 1993. Virus Genes, 2012. 44(2): p. 198–206.

64. Wong, E.H., et al., Codon usage bias and the evolution of influenza A viruses. Codon Usage Biases of Influenza Virus. BMC Evol Biol, 2010. 10: p. 253.

65. Huang, Z.X., et al., Position 34 of tRNA is a discriminative element for m5C38 modification by human DNMT2. Nucleic Acids Res, 2021. 49(22): p. 13045–13061.

66. Lecointe, F., et al., Lack of pseudouridine 38/39 in the anticodon arm of yeast cytoplasmic tRNA decreases in vivo recoding efficiency. J Biol Chem, 2002. 277(34): p. 30445–53.

67. Faivre, B., et al., Dihydrouridine synthesis in tRNAs is under reductive evolution in Mollicutes. RNA Biol, 2021. 18(12): p. 2278–2289.

68. Rappol, T., et al., tRNA expression and modification landscapes, and their dynamics during zebrafish embryo development. Nucleic Acids Res, 2024.

69. Suzuki, T., The expanding world of tRNA modifications and their disease relevance. Nat Rev Mol Cell Biol, 2021. 22(6): p. 375–392.

70. Kouno, T., et al., Crystal structure of APOBEC3A bound to single-stranded DNA reveals structural basis for cytidine deamination and specificity. Nat Commun, 2017. 8: p. 15024.

71. Fu, H., et al., Stress induces tRNA cleavage by angiogenin in mammalian cells. FEBS Lett, 2009. 583(2): p. 437–42.

72. Yamasaki, S., et al., Angiogenin cleaves tRNA and promotes stress-induced translational repression. J Cell Biol, 2009. 185(1): p. 35–42.

73. Emara, M.M., et al., Angiogenin-induced tRNA-derived stress-induced RNAs promote stress-induced stress granule assembly. J Biol Chem, 2010. 285(14): p. 10959–68.

74. Ivanov, P., et al., Angiogenin-induced tRNA fragments inhibit translation initiation. Mol Cell, 2011. 43(4): p. 613–23.

75. Saikia, M., et al., Angiogenin-cleaved tRNA halves interact with cytochrome c, protecting cells from apoptosis during osmotic stress. Mol Cell Biol, 2014. 34(13): p. 2450–63.

76. Chu, X., et al., Transfer RNAs-derived small RNAs and their application potential in multiple diseases. Front Cell Dev Biol, 2022. 10: p. 954431.

77. Pandey, K.K., et al., Regulatory roles of tRNA-derived RNA fragments in human pathophysiology. Mol Ther Nucleic Acids, 2021. 26: p. 161–173.

78. Kumar, P., C. Kuscu, and A. Dutta, Biogenesis and Function of Transfer RNA-Related Fragments (tRFs). Trends Biochem Sci, 2016. 41(8): p. 679–689.

79. George, S., et al., tRNA derived small RNAs-Small players with big roles. Front Genet, 2022. 13: p. 997780.

80. Schwartz, M.H. and T. Pan, Function and origin of mistranslation in distinct cellular contexts. Crit Rev Biochem Mol Biol, 2017. 52(2): p. 205–219.

81. Lant, J.T., et al., Genetic Interaction of tRNA-Dependent Mistranslation with Fused in Sarcoma Protein Aggregates. Genes (Basel), 2023. 14(2).

82. Netzer, N., et al., Innate immune and chemically triggered oxidative stress modifies translational fidelity. Nature, 2009. 462(7272): p. 522–6.

83. Oh, S., et al., Genotoxic stress and viral infection induce transient expression of APOBEC3A and pro-inflammatory genes through two distinct pathways. Nat Commun, 2021. 12(1): p. 4917.

84. Cortez, L.M., et al., APOBEC3A is a prominent cytidine deaminase in breast cancer. PLoS Genet, 2019. 15(12): p. e1008545.

85. Nibert, M.L., Mitovirus UGA(Trp) codon usage parallels that of host mitochondria. Virology, 2017. 507: p. 96–100.

86. Truitt, M.L. and D. Ruggero, New frontiers in translational control of the cancer genome. Nat Rev Cancer, 2016. 16(5): p. 288–304.

87. Goodarzi, H., et al., Modulated Expression of Specific tRNAs Drives Gene Expression and Cancer Progression. Cell, 2016. 165(6): p. 1416–1427.

88. Ljungstrom, V., et al., Whole-exome sequencing in relapsing chronic lymphocytic leukemia: clinical impact of recurrent RPS15 mutations. Blood, 2016. 127(8): p. 1007–16.

89. Schmutzer, M. and A. Wagner, Not Quite Lost in Translation: Mistranslation Alters Adaptive Landscape Topography and the Dynamics of Evolution. Mol Biol Evol, 2023. 40(6).

90. Pan, T., Adaptive translation as a mechanism of stress response and adaptation. Annu Rev Genet, 2013. 47: p. 121–37.

91. Bratulic, S., M. Toll-Riera, and A. Wagner, Mistranslation can enhance fitness through purging of deleterious mutations. Nat Commun, 2017. 8: p. 15410.

92. Starczak, M., et al., Quantification of DNA Modifications Using Two-Dimensional Ultraperformance Liquid Chromatography Tandem Mass Spectrometry (2D-UPLC-MS/MS). Methods Mol Biol, 2021. 2198: p. 91–108.

93. Chan, P.P. and T.M. Lowe, GtRNAdb: a database of transfer RNA genes detected in genomic sequence. Nucleic Acids Res, 2009. 37(Database issue): p. D93–7.

94. Abramson, J., et al., Accurate structure prediction of biomolecular interactions with AlphaFold 3. Nature, 2024. 630(8016): p. 493–500.

95. Bowers, K.J., et al. Scalable algorithms for molecular dynamics simulations on commodity clusters. in Proceedings of the ACM/IEEE Conference on Supercomputing (SC06). 2006. Tampa, Florida, USA.

96. Lu, C., et al., OPLS4: Improving Force Field Accuracy on Challenging Regimes of Chemical Space. J Chem Theory Comput, 2021. 17(7): p. 4291–4300.

97. Agirre, J., et al., The CCP4 suite: integrative software for macromolecular crystallography. Acta Crystallogr D Struct Biol, 2023. 79(Pt 6): p. 449–461.

98. Pettersen, E.F., et al., UCSF ChimeraX: Structure visualization for researchers, educators, and developers. Protein Sci, 2021. 30(1): p. 70–82.

