## Supplementary Information for "APOBEC3A deaminase catalyzes site-specific editing of transfer RNAs"

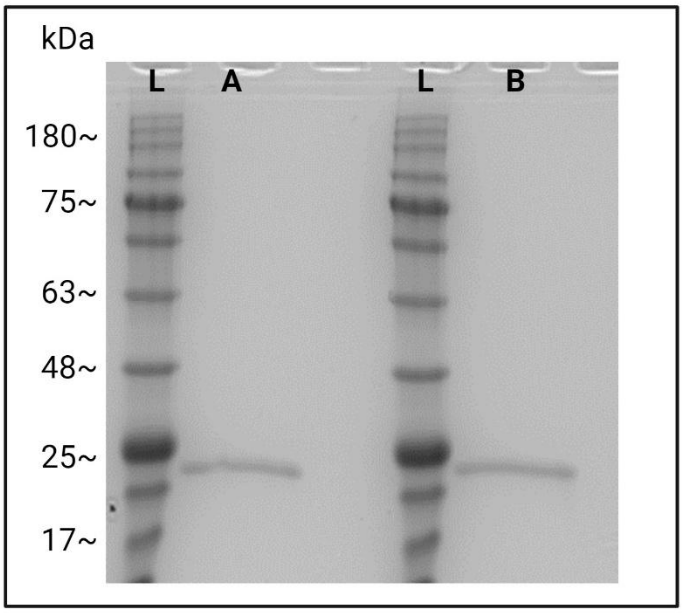

**Supplementary Figure S1.** Denaturing PAGE analysis of purified, recombinant WT_A3A (A) and E72A_A3A (B). L - Molecular weight ladder (mass is expressed in kilodaltons, kDa). Created in BioRender. Hoffa-Sobiech, K. (2025) https://BioRender.com/52fxinm.

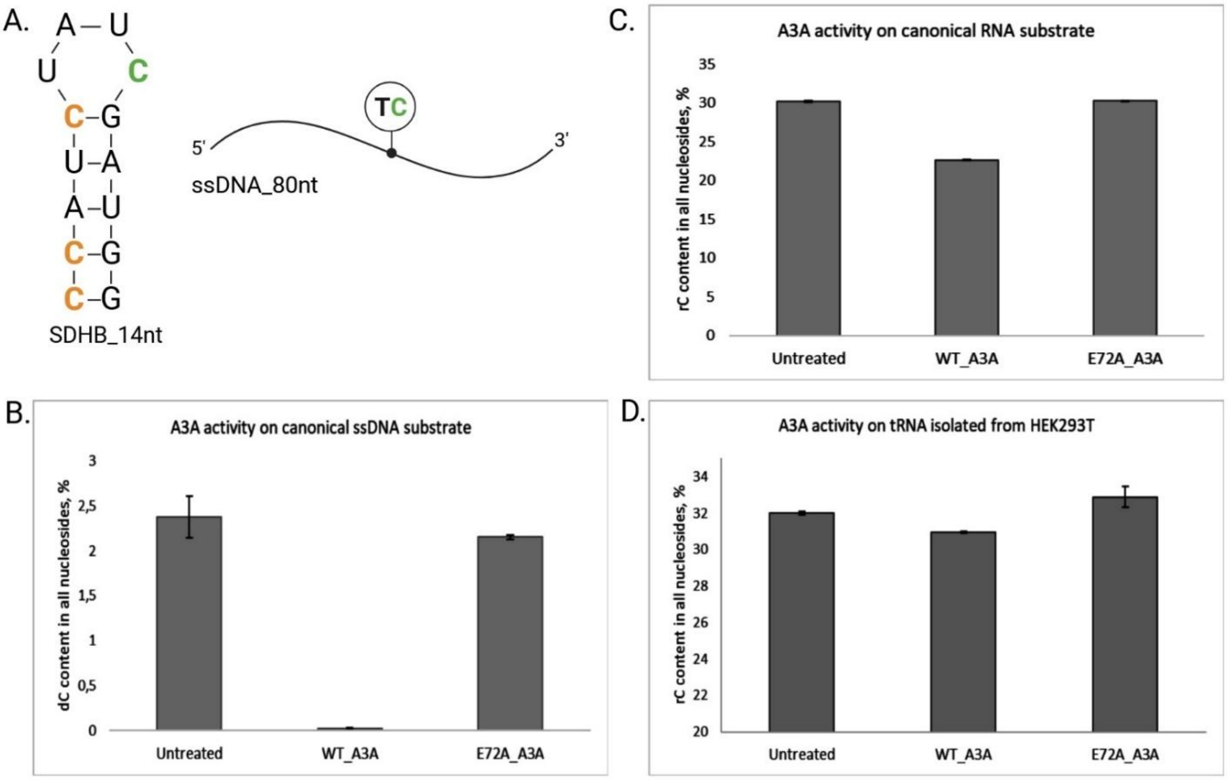

**Supplementary Figure S2. Quantitative analysis of A3A activity on canonical DNA and RNA substrates, and cell-derived tRNA.** (A) Schematic representation of the canonical ssDNA and RNA substrates. The deaminated C is shown in green. (B, C, D) A3A activity on canonical DNA (B) and RNA (C) substrates, and tRNA isolated from HEK293T cells (D). WT_A3A- or E72A_A3A-treated, and untreated substrates were digested to nucleosides and analyzed by LC-MS/MS. The deaminase activity was graphed as an average (of three replicates) dC/rC content in all nucleosides and expressed as a percentage. Created in BioRender. Hoffa-Sobiech, K. (2025) https://BioRender.com/014mwd2.

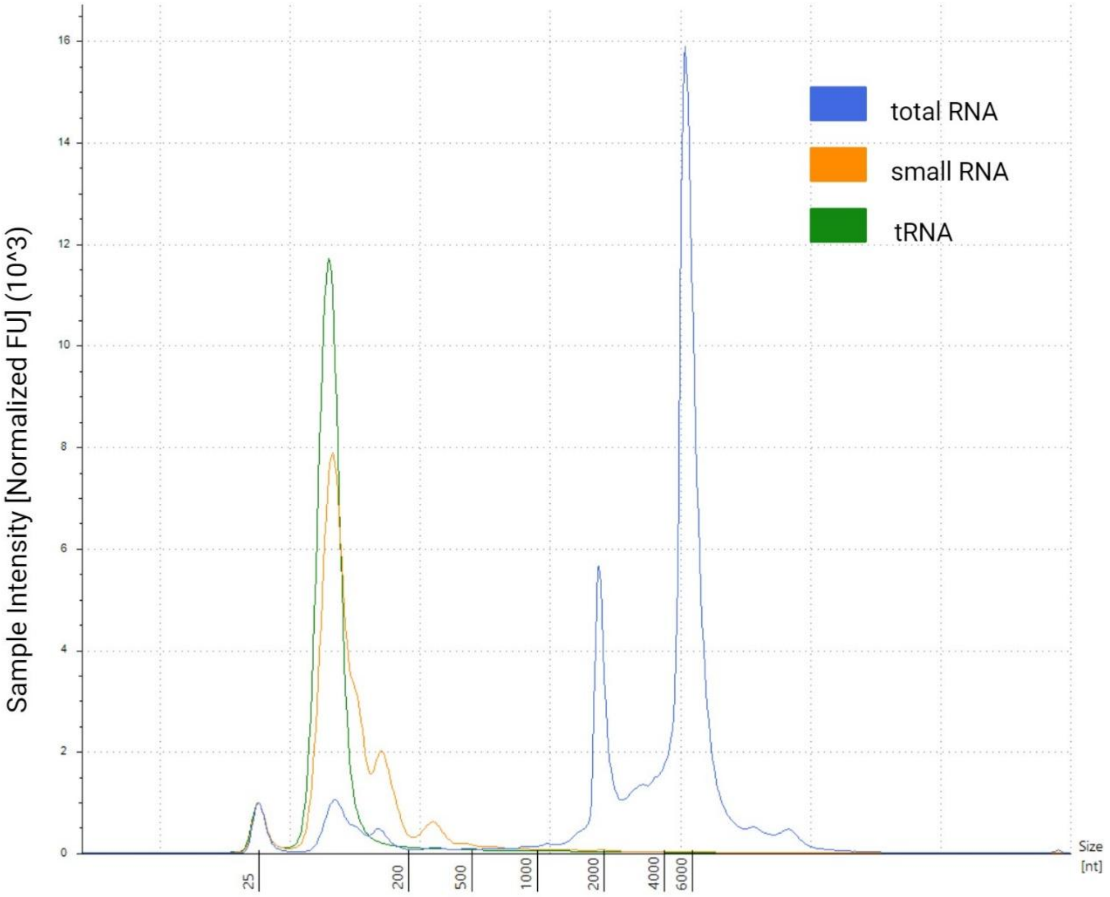

**Supplementary Figure S3.** Size distribution (in nucleotides, nt) of total RNA, small RNA, and tRNA fractions isolated from HEK293T cells. Created in BioRender. Hoffa-sobiech, K. (2025) https://BioRender.com/vkf1kja.

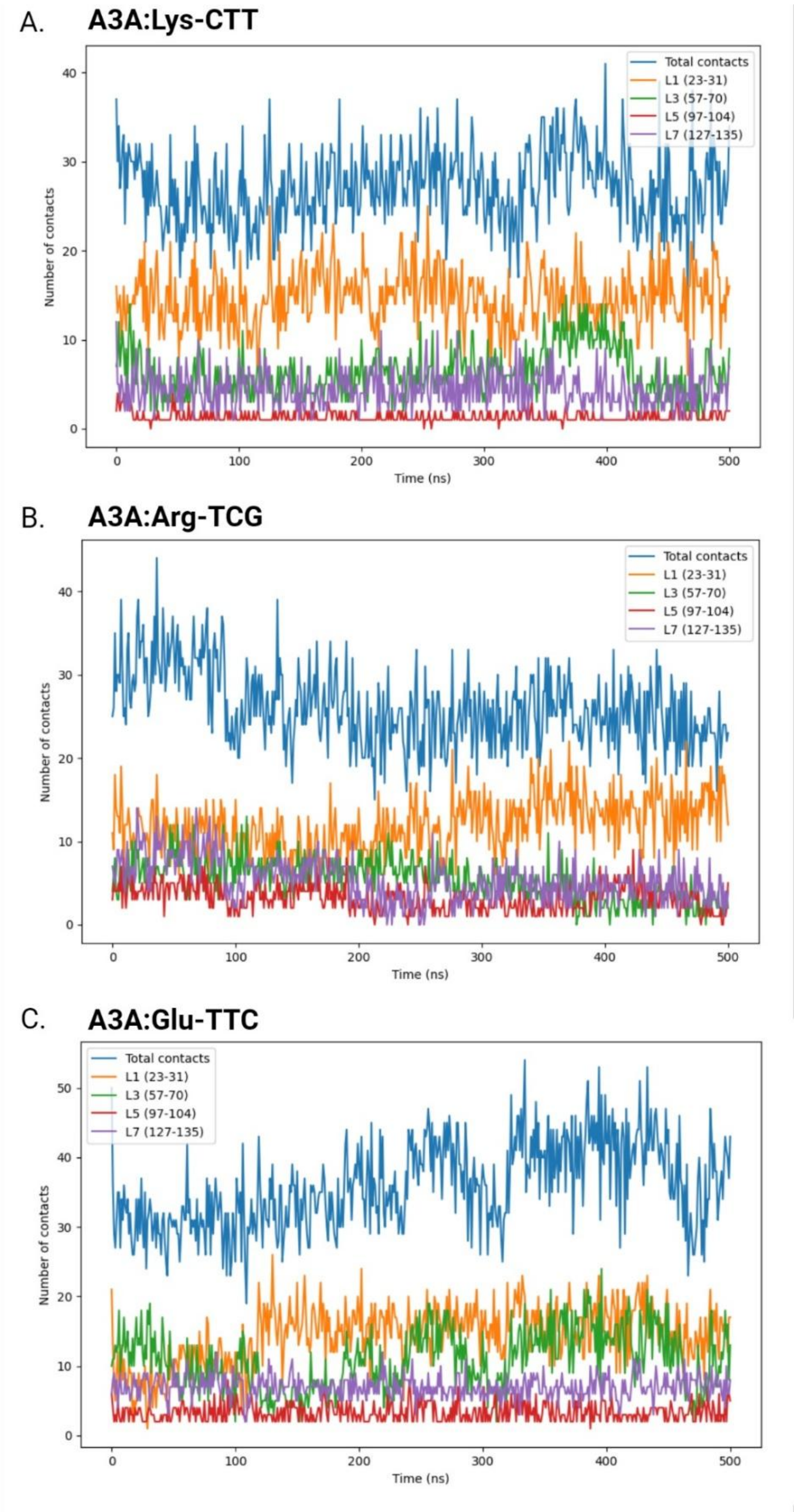

**Supplementary Figure S4.** Number of total contacts and contacts for specific loops observed throughout MD trajectories within: (A) A3A:Lys-CTT-3-1, (B) A3A:Arg-TCG-1-1, and (C) A3A:Glu-TTC-3-1 model. Color codes indicate contacts for specific loops or total contacts (see legend). The range of amino acid residues for each loop is given in brackets. Created in BioRender. Budzko, L. (2025) https://BioRender.com/hjv7pd8.

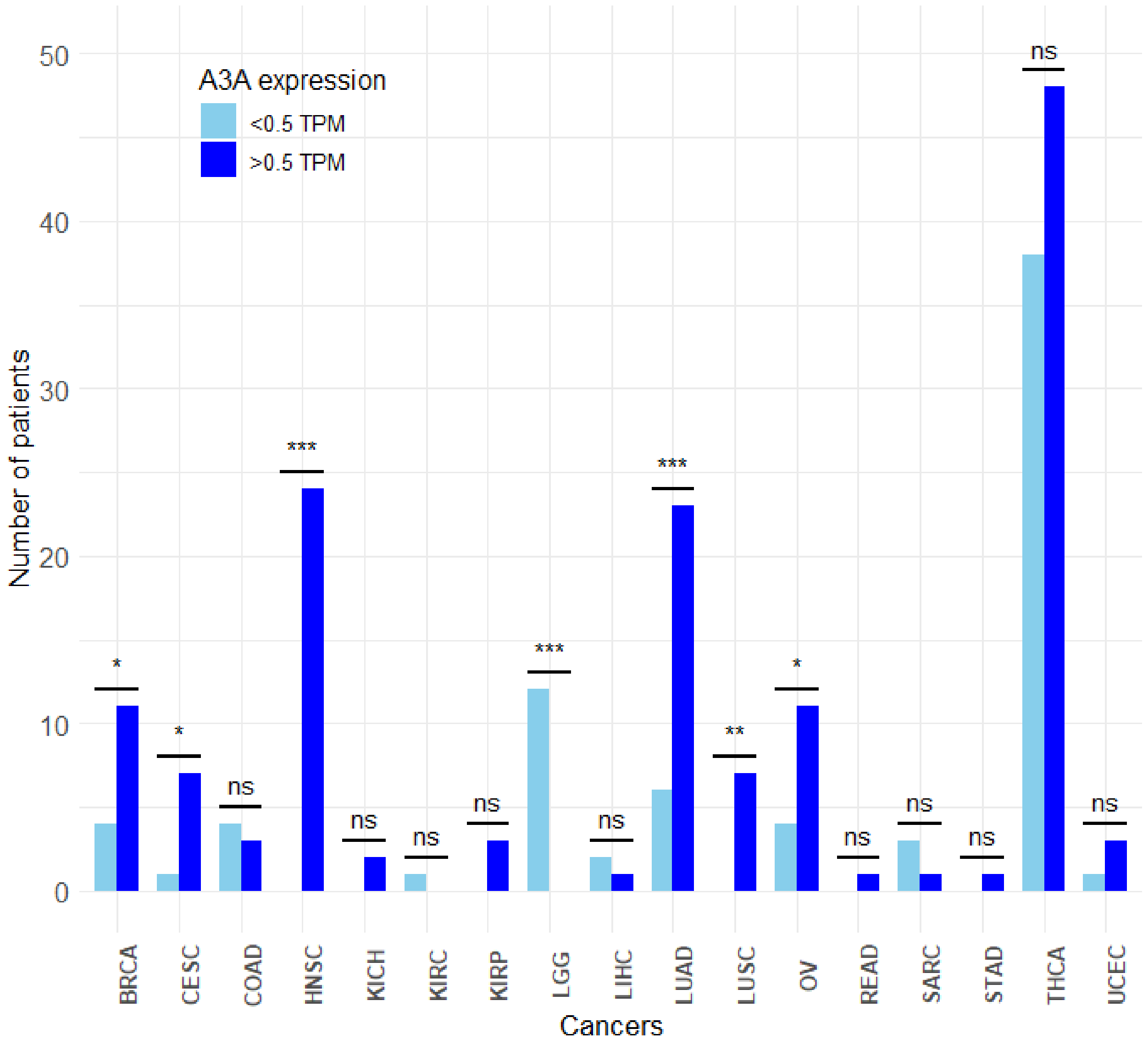

**Supplementary Figure S5.** Number of samples (in which tRF editing was detected) in individual tumor types divided by the presence (> 0.5 TPM) or absence (< 0.5 TPM) of *A3A* gene expression. The two-proportions Z test was used to assess statistical significance between these two tested groups. The figure was created in R, using the ggplot2 package.

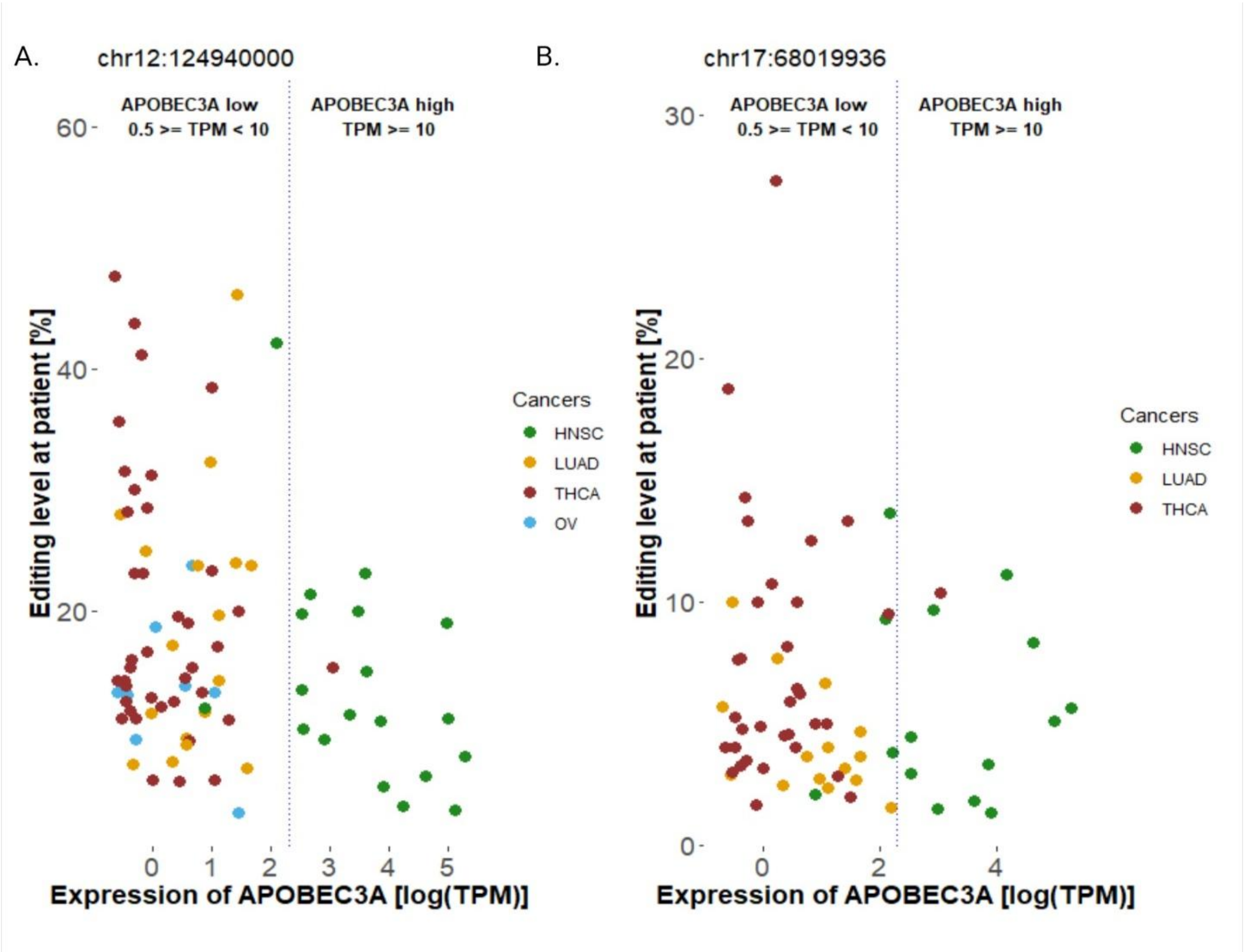

**Supplementary Figure S6.** Tumor samples plotted according to their *A3A* expression level and editing level detected in miRNA-Seq reads aligned to (A) genomic position chr12:124940000 (which corresponds to the second anticodon position in tRNA-Ala-TGC-4-1), and (B) genomic position chr17:68019936 (which corresponds to the first anticodon position in tRNA-Arg-CCG-2-1). Color codes indicate tumor types. The figure was created in R, using the ggplot2 package.

### Supplementary Methods

**Human embryonic kidney cells (HEK293T) culture**

HEK293 cells were cultured in high-glucose Dulbecco's modified Eagle's medium with sodium pyruvate (DMEM, Gibco) supplemented with 10% fetal bovine serum (FBS, Gibco) and 1% antibiotics (Penicillin/Streptomycin, Gibco). Cells were cultured in the T-75 flask at 37°C with 5% $CO_2$. Upon reaching 80 to 90% confluency, cells were dissociated using Trypsin-EDTA (0.05%) (Gibco) and passaged at a ratio of 1:5. To receive cell pellet - media was removed, cells were washed with PBS, then samples snap-frozen with liquid nitrogen. Samples were stored at -80°C until processed.

**Computational analysis of tRNA editing in NGS sequencing data**

**Data processing, mapping, and variant calling**

The single-end 150 nt reads in BLCA format, generated with the NovaSeq 6000 Illumina platform, were demultiplexed using bcl2fastq (v2.19.0) (available at: https://support.illumina.com/sequencing/sequencing_software/bcl2fastq-conversion-software.html). The raw sequencing data were trimmed with fastp (v. 0.23.2) [1] to remove the adaptor sequences and non-templated CCA additions. The quality control was performed using FastQC (v0.11.8) (available at https://www.bioinformatics.babraham.ac.uk/projects/fastqc/). The trimmed reads were excluded if the Phred Quality Score was lower than 30. Only high-quality reads 60–95 nucleotides in length were kept. The libraries were normalized by the number of reads. For each library, 40 million reads were randomly selected with seqtk (v. 1.3-r106) (available at: https://github.com/lh3/seqtk), and a selected read could be chosen only once. The filtered reads were then aligned to the hg38 human genome (downloaded from the TCGA database v.36 ), using STAR aligner (v.2.7.8a) [2]. The following STAR options were used: *--outFilterMismatchNmax 3* to filter out reads with more than 3 mismatches*; --outFilterMultimapScoreRange 0* to only report the alignments with the best scores; *--outFilterMultimapNmax 10* to discard reads that were mapped to more than 10 locations; *--alignIntronMin 1* to avoid splice alignments, *--alignEndsType EndToEnd* to force end-to-end read alignment. All reads were divided based on DNA strands and were excluded if MAPQ was lower than 20 with SAMtools (v.1.2) [3]. The above steps caused uniquely mapped reads to be selected for further analysis. The boundaries of tRNA genes were determined based on coordinates from gtRNAdb (release 21) [4]. The uniquely mapped reads were filtered out with bedtools complement (v2.27.1) [5] and SAMTools view if they exceeded the strand-specific tRNA boundaries by at least 1 bp. For further analysis, only tRNA isodecoders having a coverage 250x were used (the threshold was calculated based on the 20th percentile of tRNA coverage from each library and then calculated as an average). The variant calling was performed on strand-specific BAM files with freebayes [6] and parameters as follows: *--no-indels, --no-complex, --haplotype-length 0* to call of single-base variants that occur side by side; *--use-duplicate-reads* to include duplicate-marked alignments in the analysis, *--min-base-quality 30* to filter out variants with a base quality score lower than 30; *--min-alternate-count 3*, *--min-alternate-fraction 0.01* to evaluate positions with at least 3 reads supporting an alternate allele and they constituted at least 1% of reads.

**Identification of tRNA editing**

To investigate tRNA editing, C>T and G>A substitutions were annotated by mapping to strand-specific tRNA regions, using bedtools intersect (v2.27.1) [5]. The number of bases (A, T, C, or G) for each site was calculated by DepthOfCoverage in gatk (v. 4.2) [7] with the *--print-base-counts true* option. The C>T and G>A substitutions were considered to be false positives and were removed if: (i) the sum of A and G nucleobases were greater than T nucleobases for C>T detected sites; (ii) the sum of T and C nucleobases were greater than A nucleobases for A>G detected sites; (iii) were consistent with mismatches between RNA-central derived tRNA sequences and tRNA genes (see below); (iv) if the minimum depth at 250 reads was not detected for at least two replicates among the WT_A3A-treated samples; (v) if the minimum depth at 250 reads was not detected for all control samples. The C>T editing level was calculated as the ratio of the number of variants T to the total number of reads (C+A+T+G) at each site of detected editing on the positive strand and multiplied by 100%. The G>A editing level was calculated as the ratio of the number of variants A to the total number of reads (C+A+T+G) at each site of detected editing on the negative strand and multiplied by 100%. For each site of detected editing in WT_A3A-treated samples, a background editing level in control samples was estimated. The fold-change of detected editing between pairs of WT_A3A-treated and control samples was calculated (separately for E72A control and untreated). The detected editing sites for which the fold-change was equal or higher than 2 (in comparison to both types of controls) were used for downstream analysis. Finally, the location of the detected editing sites within the tRNA secondary structure was determined (according to tRNA secondary structures presented in gtRNAdb). To generate a genomic view of detected editing sites, the BAM files were visualized by the Integrative Genomics Viewer (IGV) [8] on the hg38 genome, downloaded from TCGA.

**Identification of mismatches between tRNA sequences and tRNA genes**

Human tRNA sequences were downloaded from RNAcentral (release 17.0) [9] database to identify mismatches between tRNA sequences and tRNA genes in the hg38 human genome (that could introduce false positive editing sites). RNAcentral-derived tRNA sequences were mapped to the hg38 human genome with bowtie (v.1.0) [10]. All valid alignments (*--all*) with a maximum of 3 mismatches (*--v 3*) on the best strand (*--best*) were reported. The SAM file was converted to the BAM format and sorted by coordinates with SAMtools. Next, freebayes (v. 1.3.8) was reused to call the variants with the changed values for the following options: *--min-coverage 1*, *--min-alternate-count* 1. C>T mismatches were selected from the positive strand and G>A from the negative strand.

**Analysis of replication probability of results**

The bootstrap analysis was performed to analyze the replication probability of the obtained results [11]. The previously normalized and restricted to 40 million reads libraries were resampled 1000 times with fastq-tools (v. 0.8.3) (available at: https://github.com/dcjones/fastq-tools) with replacement option. Next, reads were mapped and variant calling was performed, as described above. Bootstrap accuracy level and confidence interval (CI) were calculated for the primary editing sites (detected in the primary analysis). For some positions, we observed a smaller number of bootstraps reported than the expected 1000 due to a lack of minimal depth and/or editing (see Supplementary Table 2). For further analysis, we required occurrence of editing in the same replicates (among the WT_A3A-treated samples) as in the primary analysis, ensuring consistency in the number of replicates. The bootstrap accuracy level for each primary site was established as a ratio of the number of consistent bootstraps with detected editing to 1000 (resampling size) then multiplied by 100%. Next, sites consistent with primary variants were processed to calculate 95% CI from the distribution of editing levels across all bootstraps. This method [12] used the formula: $\bar{x} \pm 1.96 * (s/\sqrt{n})$ , where: *x* parameter was the mean of editing level from all bootstraps; *s* parameter was the standard deviation; and *n* was the number of consistent bootstraps. The primary editing site was considered high-confident if: (i) the resampling had a minimum 95% bootstrap accuracy; (ii) the mean editing of the primary analysis was between the lower and upper bound of bootstrap distribution within the 95% CI. The results of this analysis are presented in Supplementary Table 2.

**Computational analysis of tRF editing in TCGA data**

**Published datasets**

To validate the *in vitro* edited positions in sequencing data deposited in TCGA database (v.36) (https://www.cancer.gov/tcga), we analyzed 1230 samples (19 cancer types) for which data sets consisting of miRNA-seq, RNA-seq, and WGS data were available (Supplementary Table S3). The RNA-seq data were downloaded as an expression quantification data type to estimate the *A3A* expression level. The tumor-normal WGS data were used to call germline and somatic variants. The MINTbase database (v2.0) (http://cm.jefferson.edu/MINTbase/) (which includes also TCGA data) [13] was used as the source of tRNA coordinates for which at least one tRF was reported in the database.

**Processing the published datasets and variants calling**

The tRNA boundaries from MINTbase (with at least one tRF read) were moved from hg19 assembly to hg38 by LiftOver, downloaded from UCSC (http://genome.ucsc.edu) [14]. Next, TCGA data were processed as follows. BAM files containing miRNA-seq alignments were sorted with SAMtools with -n option to sort files by read name. Next files were converted to FASTQ format by bamtofastq function in bedtools. The miRNA-Seq reads were trimmed with fastp using the following options: --*qualified_quality_phred 20 to* exclude reads with the phred quality score lower than 20, --*unqualified_percent_limit 40 to* discard *reads* if more than 40% of bases in a read had a quality score below 20, --*cut_front* to trim low-quality bases from 5` end, --*n_base_limit 5* to remove reads with at the least 5 of ambiguous (N) bases, --*length_required 15* to ensure a minimum read length at 15 nucleobases. Trimmed reads were mapped against the reference genome using STAR aligner (v.2.7.8a) as previous data but with extra options: --*outFilterMatchNminOverLread 0*, --*outFilterScoreMinOverLread* 0 to remove any limits on the mapped length, --*outFilterMatchNmin 16* to filter alignments if the number of matched bases were ≥16. The aligned reads were divided based on DNA strands and were kept if MAPQ was more than 20 by SAMTools. Next, aligned reads that exceeded tRNA boundaries by at least 1 bp were removed, using strand-specific tRNA coordinates from MINTbase. The variant calling in miRNA-Seq data was performed using freebayes as previously for *in vitro* data, with changed values of parameters as follows: --*min-alternate-count* 2, --*min-coverage* 10. Next, WGS data were processed to call somatic and germinal variants. The WGS reads in BAM file format were divided based on DNA strands and were excluded if MAPQ was lower than 20 by SAMtools. The somatic and germinal mutations were called with Mutec2 of gatk [7], using the following options: --*min-base-quality-score 15* to exclude bases with a Phred score lower than 15; --*callable-depth 1* to only consider positions with at least 1 read for variant calling, --*genotype-germline-sites true* to genotype both germline and somatic variants.

**Identification of tRF editing**

The list of high-confident *in vitro* edited sites shown in Supplementary Table 2 was used to verify they overlap with the results of variant calling in miRNA-seq TCGA data. The consistent positions were considered to be edited if they met the following criteria: (i) the editing level was ≥ 1%; (ii) the depth of specific position was at least 10 in both the miRNA-Seq and genomic tumor-normal datasets; (iii) in the miRNA-Seq data the sum of A and G nucleobases were greater than T nucleobases for C>T detected sites or sum of T and C nucleobases were greater than A nucleobases for A>G detected sites; and (iv) lack of somatic mutation and/or somatic-germline variant detected at this position in all patients. The A, T, C, or G counts at each site were calculated with the DepthOfCoverage function in gatk [7].

**Statistical analysis of tRNA editing level with A3A expression**

For samples with detected editing, RNA-Seq data downloaded from TCGA were used to estimate the *A3A* expression level. The FPKM values were converted to TPM with fpkm2tpm function in R. The *A3A* gene was considered to be expressed if TPM value was > 0.5 (denoted A3A+ samples; samples without detectable *A3A* expression, < 0.5 TPM, were denoted A3A-). To assess whether the occurrence of the editing phenomenon was more frequent in A3A+ samples compared to A3A- samples, the number of A3A+ samples and A3A- samples was compared within (i) each cancer type and (ii) across different cancer types. To assess whether the proportions in the two groups were significantly different, we used the two-proportions Z test, using prop.test function in R and considering p-values <0.05 as statistically significant. Next, the editing efficiency (at positions edited in more than 100 samples) was correlated with *A3A* expression level within each cancer type. To verify the correlation, the Person correlation test was performed (considering p-values <0.05 as statistically significant) using R with stats, magrittr and dplyr packages. The *A3A* expression level and editing level according to cancer types were plotted in R using ggplot2.

**Supplementary Tables S1-S6, S8 (provided in separate Excel files)**

**Supplementary Table S1.** Nucleus-encoded tRNA isodecoders identified in WT_A3A-treated samples and controls (with technical triplicates)

**Supplementary Table S2.** tRNA editing efficiency within the identified tRNA isodecoders (C>T and G>A mutations are presented in separate sheets)

**Supplementary Table S3.** The analyzed dataset downloaded from The Cancer Genome Atlas database

**Supplementary Table S4.** tRF editing detected in TCGA samples

**Supplementary Table S5.** Editing level for the analyzed positions for each sample compared with A3A expression level

**Supplementary Table S6.** tRF editing in specific cancer types and its correlation with the A3A gene expression level

**Supplementary Table S8.** Number of raw reads obtained from libraries prepared according to the YAMAT-seq protocol

**Supplementary Table S7.** List of oligonucleotides and synthetic tRNAs used in the manuscript

| Oligo name | Nucleic acid type | Sequence 5′-3′ |
|---|---|---|
| ssDNA_80nt | DNA | GGATTGGTTGGTTATTTGTTTAAGGAAGGTGGATTAAAATCT TAATAAGGTGATGGAAGTTATGTTTGGTAGATTGATGG |
| SDHB_14nt | RNA | CCAUCUAUCGAUGG |
| tRNA-Asp-GTC-2-1 | RNA | UCCUCGUUAGUAUAGUGGUGAGUAUCCCCGCCUGUCACG CGGGAGACCGGGGUUCGAUUCCCCGACGGGGAGCCA |
| tRNA-Gly-GCC-2-1 | RNA | GCAUUGGUGGUUCAGUGGUAGAAUUCUCGCCUGCCACGC GGGAGGCCCGGGUUCGAUUCCCGGCCAAUGCACCA |

#### References

1. Chen, S., *Ultrafast one-pass FASTQ data preprocessing, quality control, and deduplication using fastp.* Imeta, 2023. **2**(2): p. e107.
2. Dobin, A., et al., *STAR: ultrafast universal RNA-seq aligner.* Bioinformatics, 2013. **29**(1): p. 15-21.
3. Danecek, P., et al., *Twelve years of SAMtools and BCFtools.* Gigascience, 2021. **10**(2).
4. Chan, P.P. and T.M. Lowe, *GtRNAdb 2.0: an expanded database of transfer RNA genes identified in complete and draft genomes.* Nucleic Acids Res, 2016. **44**(D1): p. D184-9.
5. Quinlan, A.R. and I.M. Hall, *BEDTools: a flexible suite of utilities for comparing genomic features.* Bioinformatics, 2010. **26**(6): p. 841-2.
6. Garrison E, M.G., *Haplotype-based variant detection from short-read sequencing.* arXiv, 2012. **1207.3907**.
7. McKenna, A., et al., *The Genome Analysis Toolkit: a MapReduce framework for analyzing next-generation DNA sequencing data.* Genome Res, 2010. **20**(9): p. 1297-303.
8. Thorvaldsdottir, H., J.T. Robinson, and J.P. Mesirov, *Integrative Genomics Viewer (IGV): high-performance genomics data visualization and exploration.* Brief Bioinform, 2013. **14**(2): p. 178-92.
9. The, R.C., *RNAcentral: a hub of information for non-coding RNA sequences.* Nucleic Acids Res, 2019. **47**(D1): p. D1250-D1251.
10. Langmead, B., et al., *Ultrafast and memory-efficient alignment of short DNA sequences to the human genome.* Genome Biol, 2009. **10**(3): p. R25.
11. Bogdanovic, O. and M. Vermeulen, *Correction to: TET Proteins and DNA Demethylation.* Methods Mol Biol, 2021. **2272**: p. C1.
12. Banjanovic, E.S., Osborne, J. W., *Confidence Intervals for Effect Sizes: Applying Bootstrap Resampling.* Practical Assessment, Research, and Evaluation, 2016. **21(1): 5**.
13. Pliatsika, V., et al., *MINTbase v2.0: a comprehensive database for tRNA-derived fragments that includes nuclear and mitochondrial fragments from all The Cancer Genome Atlas projects.* Nucleic Acids Res, 2018. **46**(D1): p. D152-D159.
14. Perez, G., et al., *The UCSC Genome Browser database: 2025 update.* Nucleic Acids Res, 2024.
